# How strong are correlations in strongly recurrent neuronal networks?

**DOI:** 10.1101/274480

**Authors:** Ran Darshan, Carl van Vreeswijk, David Hansel

## Abstract

Cross-correlations in the activity in neural networks are commonly used to characterize their dynamical states and their anatomical and functional organizations. Yet, how these latter network features affect the spatiotemporal structure of the correlations in recurrent networks is not fully understood. Here, we develop a general theory for the emergence of correlated neuronal activity from the dynamics in strongly recurrent networks consisting of several populations of binary neurons. We apply this theory to the case in which the connectivity depends on the anatomical or functional distance between the neurons. We establish the architectural conditions under which the system settles into a dynamical state where correlations are strong, highly robust and spatially modulated. We show that such strong correlations arise if the network exhibits an *effective* feedforward structure. We establish how this feedforward structure determines the way correlations scale with the network size and the degree of the connectivity. In networks lacking an effective feedforward structure correlations are extremely small and only weakly depend on the number of connections per neuron. Our work shows how strong correlations can be consistent with highly irregular activity in recurrent networks, two key features of neuronal dynamics in the central nervous system.

## I. INTRODUCTION

Two-point correlations are commonly used to characterize collective dynamics in extended systems [1–7]. Recent technical advances [8–10] make it possible to simultaneously record the activity of many neurons in networks in the brain. This allows for the measurement of two-point correlations for large numbers of neuronal pairs in spontaneous activity as well as upon sensory stimulation or in controlled behavioral conditions.

Correlations in neuronal activity impact the ability of networks to encode information [11–14]. Correlations are also functionally important in performing sensory, motor, or cognitive tasks [15]. For instance, correlated oscillatory activity has been hypothesized to be involved in visual perception [16]. In a recent study combining modeling, electrophysiology and analysis of behavior, we argued that non-oscillatory correlated neuronal activity in the central nervous system is essential in the generation of exploratory behavior [17]. Correlations are also important for the self-organization of neuronal networks through activity dependent plasticity [18]. Indeed, changes in synaptic strength are thought to depend on the temporal correlation of the activty of the pre and postsynaptic neurons [19, 20].

Correlation strengths depend on the brain area, [21, 22], the layer in cortex [23], stimulus conditions, behavioral states [24] and experience [25–27]. A wide range of values for correlation coefficients, from negligible [21, 28] to substantial [11, 29–33] have been reported in the last two decades. Correlation coefficients are usually higher for close-by neurons than for neurons that are far apart [31, 34–37]. In cortex, they drop significantly over distances of 200 *-* 400 *μm* [35, 37]. Recent works have reported correlations varying non-monotonically with distance [38] or correlations which are positive for close-by neurons but negative for neurons farther apart [39]. Correlation coefficients also depend on functional properties of the neurons. Neurons which code for similar features of sensory stimuli are more correlated [21, 31, 34, 37, 40].

Neurons in cortex receive recurrent inputs from several hundreds [41] to a few thousands [42] of other neurons. Individual connections can induce post-synaptic potentials in a range of 0.1 mV to several mV [35]. Thus, 50 simultaneous inputs are sufficient to trigger or suppress a spike in a neuron. These facts have been incorporated in model networks with strongly recurrent connectivity in which connection strengths are 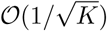, where *K* is the average number of inputs per neuron. This scaling is in contrast to the one used in standard mean-field models (e.g. [43]) where connections are *𝒪* (1*/K*) and thus are much weaker. Recent expe<u>ri</u>ments in cortical cultures are consistent with a 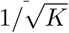 scaling of connection strength [44].

van Vreeswijk and Sompolinsky [45, 46] showed that strongly recurrent networks consisting of two populations of neurons, one excitatory (E) and one inhibitory (I), randomly connected on a directed Erdös-Ŕényi graph, operate in a state in which the strong excitation is balanced by the strong inhibition. In this *balanced regime*, neurons re<u>ce</u>ive strong excitatory and inhibitory inputs, each 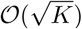, but due to the recurrent dynamics these inputs cancel each other at the leading order. This cancellation, which does not require fine-tuning, results in 𝒪 (1) net inputs into the neurons whereas spatial heterogeneities and temporal fluctuations in the inputs are also 𝒪 (1). As a result, the activity of the neurons remains finite and exhibits strong temporal irregularity and heterogeneity [47, 48].

The latter results were established in two-population sparsely connected networks. Renart et al. [28] showed that they also hold for densely connected networks. This is because in unstructured and strongly recurrent networks the dynamics suppress correlations [49]. Despite the fact that in these networks neurons share a finite fraction of their inputs, they operate in an asynchronous state with 𝒪 (1*/N*) correlations, where *N* is the number of neurons in the network [50].

These previous studies focused on strongly recurrent unstructured networks with two neuronal populations. In the brain, however, neural networks comprise a diversity of excitatory and inhibitory cell types, which differ in their morphology, molecular signature and importantly for our purpose, in their connectivity. These networks are also structured at many levels. In particular, the probability of connection falls off with distance and depends on functional properties of preand postsynaptic neurons. For example, in mouse primary auditory cortex the probability of excitatory neurons to be connected decays to zero within *∼* 300*μm* [35]. In cat primary visual cortex neurons interact locally on range of *∼* 500*μm*, whereas long range patchy connections are observed up to several *mms* [51, 52].

In the present paper we investigate how *structure* in network connectivity can give rise to strongly correlated activity. Our goal is to explore the general architectural features that control the strength of pair-wise correlations in strongly recurrent neural circuits consisting of several excitatory and inhibitory neuronal populations. In Section II, we define the network architectures and the neuronal model we use. In Section III, we establish a set of constraints, *the balanced correlation equations*, that need to be satisfied in any strongly recurrent network if firing rates do not saturate. We derive in Section IV explicit expressions for the Fourier components of the correlations in two-population networks with spatially modulated connectivity. We establish the conditions under which correlations are strong and show that these do not violate the balanced correlation equations. Section V is devoted to networks with an arbitrary number of populations. We prove two theorems, which state for which network architectures correlations are 𝒪 (1*/N*) and for which they increase with *K*, when *K* is large. In Section VI, we apply these theorems to specific examples. In Section VII, we assume that *K* = 𝒪 (*N*^*γ*^) and derive a bound on *γ* for which the scaling theorems still apply. The paper closes with a discussion of our results.

## II. THE NETWORK MODEL

### II.1. Architecture

We consider a neuronal network with a ring architecture, comprising *D* neuronal populations, some excitatory and others inhibitory (Fig. 1; [53, 54]). For simplicity, we assume that all populations have the same number of neurons, *N*. Neuron *i*, in population *α* (neuron (i,*α*)) is located at angle 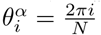 with, *i* = 1, *…, N* and *α* = 1, *…, D*. The probability, 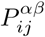, that a neuron (*j, β*) projects to neuron (*i, α*) depends on their distance on the ring (Fig. 1b). We write: 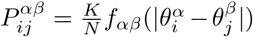 where 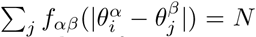. In this paper we assume a finite number of non-zero Fourier modes in *f*_*αβ*_.

**FIG. 1.**
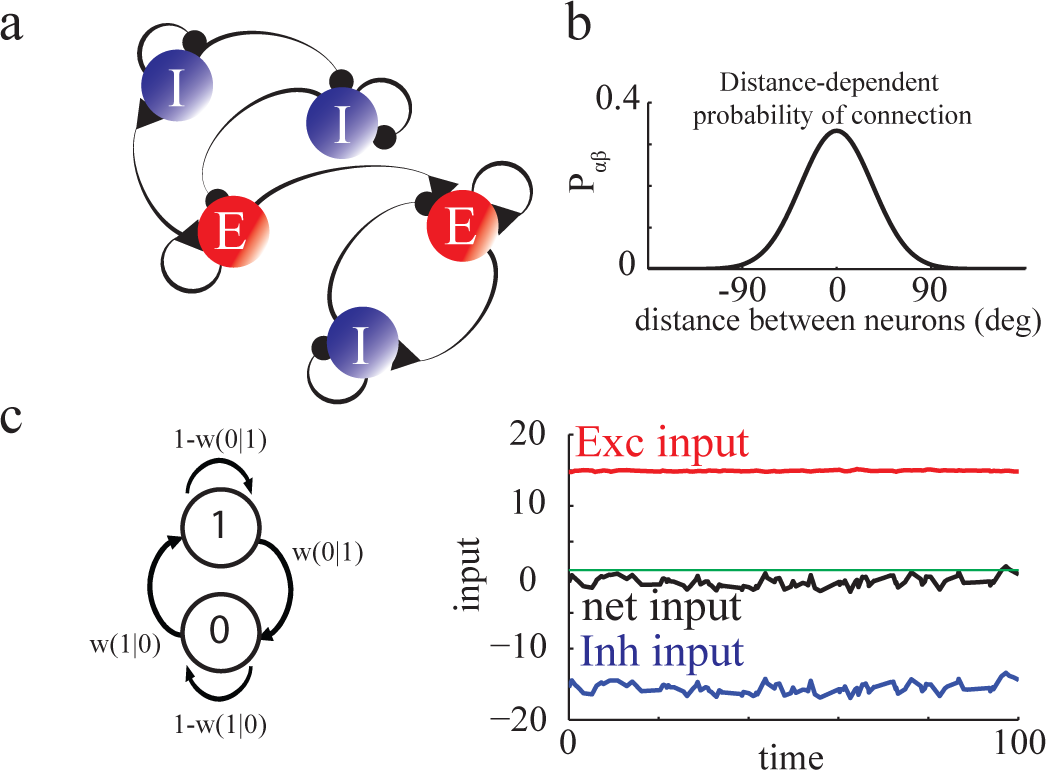
Network architectures and dynamics. **a.** Networks consist of *D* populations of neurons, excitatory (*E*,red) and inhibitory (*I*, blue), recurrently coupled and receiving external drives. Triangles: Excitatory connections. Circles: Inhibitory connections. **b.** The neurons in each population are located on a ring. The probability that neuron (*j, β*) is connected to neuron (*i, α*) depends on their distance. **c.** Left: Neurons are binary units with zero temperature Glauber dynamics. Right: The network operates in the balanced regime. Red: Excitatory input into a neuron. Blue: Inhibitory input into the same neuron. Black: The net input is on the order of the threshold (green, *T* = 1). Time is given in units of *τ* = 1.

Thus, a neuron in population *α* receives, on average, *K* inputs from neurons in each of the populations *β*. We denote by **λ** the adjacency matrix of the network connectivity

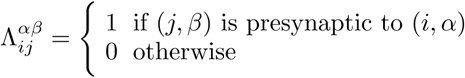

For simplicity we assume that all connections from population *β* to population *α* have the same strength, *j*_*αβ*_. We thus define the connectivity matrix, ***J***, as

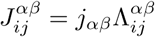

Note that *j*_*αβ*_ is positive (negative) if population *β* is excitatory (inhibitory).

In all this paper we focus on strongly recurrent networks, characterized by interactions which are 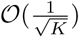 [45, 46]. Thus, we scale the synaptic strengths with the mean connectivity as

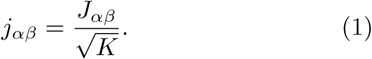

where *J*_*αβ*_ is 𝒪 (1).

### II.2. Single neuron dynamics

The state of neuron (*i, α*) is characterized by a binary variable, 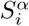. When the neuron is quiescent,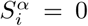 while if it is active, 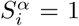. The total synaptic current into neuron (*i, α*) at time *t*, 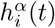, is the result of all its recurrent interactions with the other neurons in the network, as well as the feedforward inputs coming from outside the network. It is given by

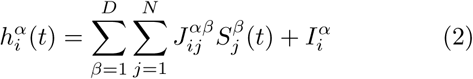

where the external input 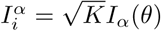 [46] is assumed to be constant in time. Here, *I*_*α*_ is 𝒪 (1) and thus 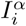 is 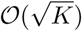.

The neurons are updated with Glauber dynamics at zero temperature [1, 28, 43]. Specifically, update times for neuron (*i, α*) are Poisson distributed with rate 1*/τ*: if neuron (*i, α*) is updated at time *t*, 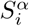 is set to 1 if 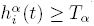 and to 0 otherwise (for simplicity, we assume the same threshold, *T*_*α*_, for all neurons in population *α*, and the same update rate for all neurons). Accordingly, the transition probability, *w*, (Fig. 1c) can be written as

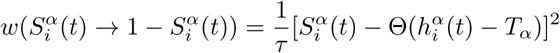

where Θ is the Heaviside function. We normalize time so that *τ* = 1.

## III. SPATIO-TEMPORAL PROFILE AND CORRELATIONS OF THE ACTIVITY FOR *N, K → ∞*

We write the state of neuron (*i, α*) as

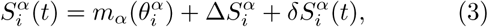

Where 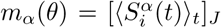 is the average of 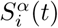 over realizations of the network connectivity matrix and over time. In Eq. (3), the term 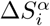 is the quenched disorder, 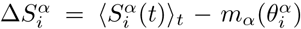, whereas 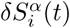 represents the temporal fluctuations in the activity of neuron (*i, α*),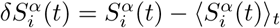.

Similarly, we can write:

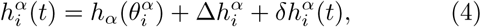

Where 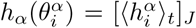 is a smooth function of its argument. For large *N* it is given by:

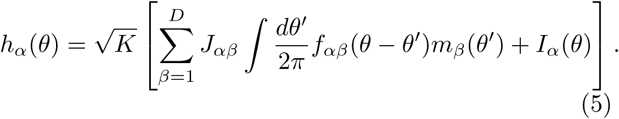

The second term in Eq. (4) is the quenched disorder in the input. It satisfies

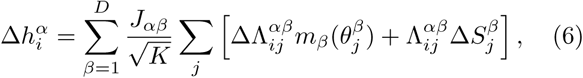

where 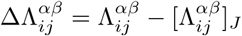.

Finally, the temporal fluctuations in the inputs are

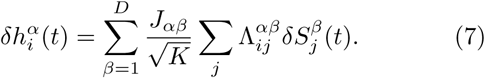

Because we scale the synaptic strength as 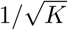, these temporal fluctuations are 𝒪 (1) <u>wh</u>en correlations are weak. At first sight,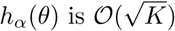. This would imply that, depending on whether *h*_*α*_(*θ*) is positive or negative, the neurons fire either at very high or at very low rate. This happens unless the network settles into a state in which excitation and inhibition are, on average, balanced. In this case, the net average inputs to the neurons are in fact 𝒪 (1). On the other hand, large spatial and temporal correlations could in principle lead to temp<u>or</u>al fluctuations and heterogeneities in the inputs of 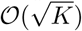. Thus in presence of strong correlations it is not sufficient to require that the mean input, *h*_*α*_(*θ*), is 𝒪 (1), for the network to operate in a biologically relevant regime. For that, we need both the mean net inputs and the input fluctuations to be 𝒪 (1) at any time and for all neurons. When this happens we say that the network operates in the balanced regime.

### III.1 Balance equations for the quenched averaged population activities

As in [55, 56], the requirement that the mean input is 𝒪 (1) yields

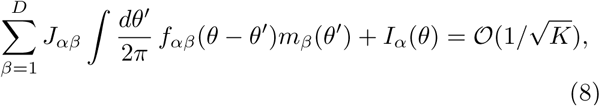

for all *α ∈*{1, *…, D*} and all *θ ∈* [0, 2*π*). In the large *K* limit this yields a set of linear equations which determines the functions *m*_*α*_(*θ*) to leading order. These equations can be written in Fourier space

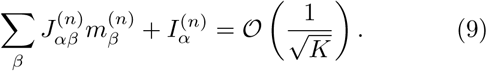

where the superscript *n* denotes the *n*^*th*^ Fourier mode and we have used the short hand notation

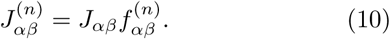

which is the *n*^*th*^ Fourier mode of the connectivity matrix. In what follows, we consider 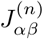 as the elements of a *D × D* matrix, ***J***^(n)^.

The spatial average of the activities, 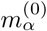, must be non-negative for the balanced state to exist. This implies that the parameters *J*_*αβ*_ and the external inputs,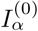, must satisfy a set of inequalities. For example, for networks with two populations, one excitatory (*α* = *E*) and one inhibitory (*α* = *I*), these inequalities are [46]

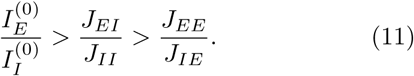

In general, Eq. (9), also implies additional inequalities that must be satisfied by 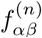 to guarantee that *m*_*α*_(*θ*) *≥* 0 for all *α* and *θ* [55, 56]. In the present work we focus on the case where the external inputs are spatially homogeneous. Thus,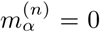 for *n ≥* 1, and the only condition which is required for the balanced state to exist is: 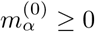.

To study the stability of the homogeneous balanced state it is useful to introduce the *interaction* matrix

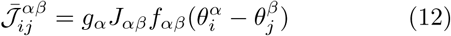

where *g*_*α*_ is the population averaged gain (see Appendixes A,D; [46, 49]).

Small perturbations, *δm*_*α*_(*θ, t*), of the activity profile around this homogeneous state evolve according to [46]:

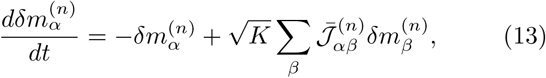

where 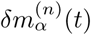 and 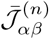are the *n*^*th*^ Fourier modes of *δm*_*α*_(*θ, t*) and 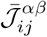. Since each row of 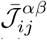 has 𝒪 (*K*) non-zero elements which are 𝒪(1*/K*),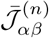 is 𝒪 (1).

The stability of the balanced state with respect to perturbations in *m*_*α*_(*θ*) requires that, for all *n*, all the eigenvalues of the matrices ***𝒯***^(n)^have real parts smaller than 1. For instance, for a two population (E,I) network one must have in the large *K* limit

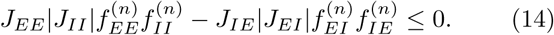

for all *n*. Note that the gain of the neurons, *g*_*α*_, dropped from this equation. The balanced state undergoes a Turing bifurcation when for some 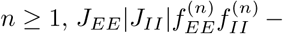 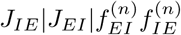 crosses 0 and becomes positive.

### III.2. Balance equation for pair-wise correlations

The time-lagged auto- and cross-correlation functions of the activities of a pair of neurons, (*i, α*), (*j, β*), for (*j, β*) ≠ (*i, α*), are

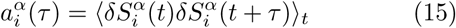

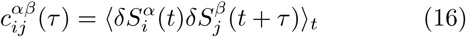

In what follows, it is convenient to define 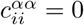.

Due to the randomness of the connectivity, the number of excitatory and inhibitory inputs varies from neuron to neuron, resulting in heterogeneous firing rates between neurons even when they belong to the same population [46, 48]. This structural randomness also results in heterogeneity of the auto-correlations of single neuron activities. It also contributes to the heterogeneity in pair cross-correlations. The latter is further enhanced by the spatial variability in the number of inputs shared by pair of neurons. A full characterization of the distributions of the auto- and cross-correlations is beyond the scope of this paper. Instead, here we will focus on their population averages.

The population average auto-correlation, *A*_*α*_, is given by 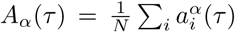, which in the thermodynamic limit is also 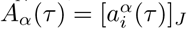. With the architecture we use, the probability that neurons (*i, α*) and (*j, β*) share common inputs from a third neuron, (*k, γ*) is

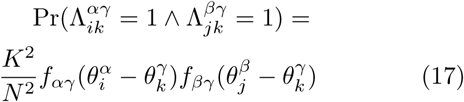

As is evident from this equation, the number of shared inputs averaged over all pairs of neurons separated by the same distance on the ring, Δ, depends only on Δ. Thus we define the average cross-correlations as 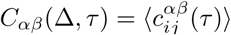, where ⟨. ⟩ denotes the average over pairs with 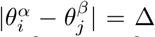. In the thermodynamic limit, this quantity does not depend on the specific realization of the network connectivity matrix. The Fourier expansion of this function is

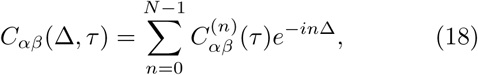

where 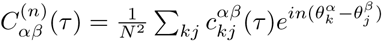.

Equation (9), which determines the population averaged firing rates, stems from the constraint that the net input in every neuron must be 𝒪 (1) when *K* is large. The condition that for any pair of neurons the correlation in their inputs is finite in that limit leads to another constraint, which we now derive.

Let us consider the *ND × ND* matrix ***Q*** defined by

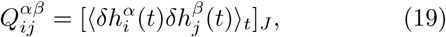

for 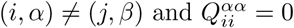. Using Eq.(7), one finds

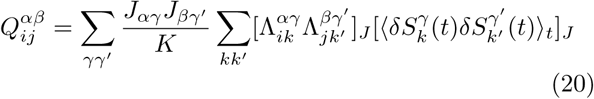

for (*i, α*) ≠ (*j, β*). Similarly to *C*_*αβ*_(Δ), 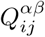 is a function of 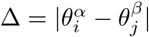 only.

Expanding Eq.(20) in Fourier 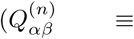 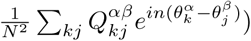, yields

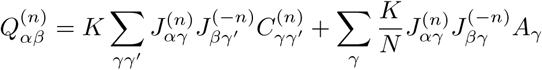

which can be written in matrix form

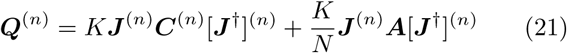

Here ***X*** denotes the *D × D* matrix *X*_*αβ*_, ***A***_*αβ*_ = *A*_*α*_*δ*_*αβ*_ and the superscript *†* denotes Hermitian conjugation 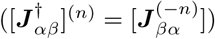.

Note that the diagonal of the matrix ***Q*** is not the variance of the inputs. The latter is (see Appendix E4)

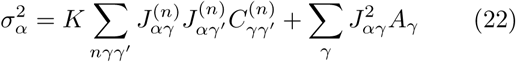

The requirement that crosscorrelations of the inputs into pair of neurons are at most 𝒪 (1) implies that all the quantities 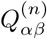 are also at most 𝒪 (1). This yields

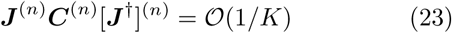

We call Eq. (23), the *balanced correlation equation*. It implies that for all *n* for which the matrix ***J***^(n)^is invertible and has entries 𝒪 (1), ***C***^(n)^is smaller or equal to 𝒪 (1*/K*) and is thus weak. For a broad class of network architectures, we show below that correlations are in fact 𝒪 (1*/N*) and barely depend on *K*, for large *K*. When, however, ***J***^(n)^is singular for some *n ≥* 0, correlations can be larger than 𝒪 (1*/K*). In fact we will see that in those cases correlations can even increase with *K*.

Finally, we note that one can also write a balanced correlation equation for the quenched disorder of the neural activity, using the fact that 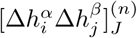 also must be at most 𝒪 (1). This requirement leads to the condition

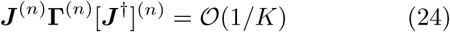

With

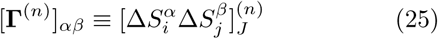

## IV. CORRELATIONS IN TWO-POPULATION NETWORKS

In Appendix A we derive an equation for the spatial Fourier modes of the equal-time quenched average correlation functions, *C*_*αβ*_(Δ, 0). It yields (omitting the second argument)

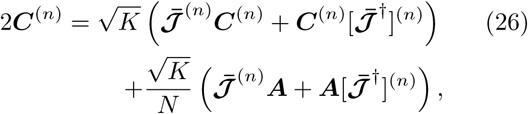

There we also show that the solution of this equation is a fixed point of the dynamics of the correlations which is always stable when the balanced state is stable with respect to perturbation in the population rates.

This equation holds for a network with an arbitrary number of neuronal populations. In the case of two populations, it can be solved explicitly, yielding after a straightforward calculation (see Appendix E)

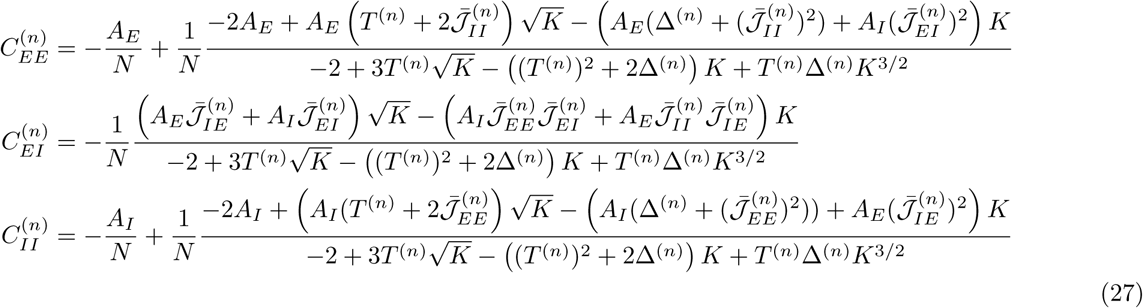

where *T* ^(n)^and Δ^(n)^are the trace and the determinant of the matrix 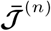.

The expansion of these expressions for large *K* gives

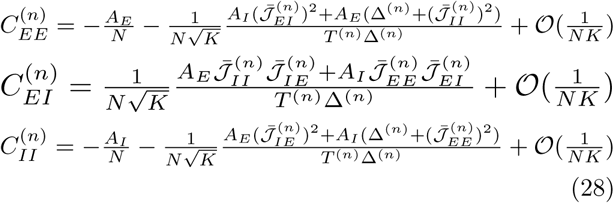

Thus in general when *K* is large 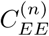 and 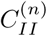 are very small, namely 𝒪 (1*/N*), with negative prefactors which do not depend on *K*. As for 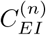, it is smaller than 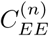 and 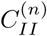 by a factor 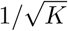.

It should be noted that to derive Eq. (28) we assumed that *T* ^(n)^≠0 and Δ^(n)^≠0. Equation (27) indicates, however, that when *T* ^(n)^= 0 and Δ^(n)^≠ 0, or *T* ^(n)^≠ 0 and Δ^(n)^= 0,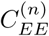,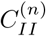 and 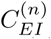 are 𝒪 (1*/N*).

The situation is different if *T* ^(n)^= 0 *and* Δ^(n)^= 0. Equation (27) shows that in this case it is possible to get correlations which are 𝒪 (*K/N*).

In the rest of this section we consider in detail two-population networks in which for the probabilities of connection only the first two Fourier modes are non-zero

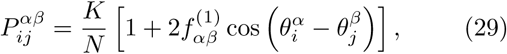

with *α* = *E, I, β* = *E, I*.

### IV.1. *T*^(^1^)^ ≠0 and Δ ^(^1^)^ ≠ 0

For 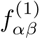 such that *T* ^(^1^)^ ≠ 0 and Δ^(^1^)^ ≠ 0 we have in the large *N, K* limit

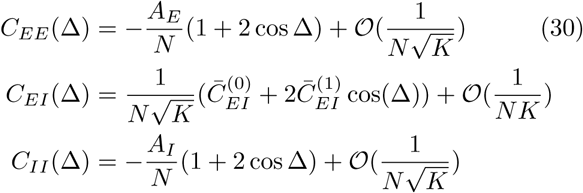

Where 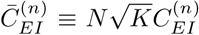 are 𝒪 (1). Thus, in that limit, the spatial average and modulation of the correlations within the E and I populations do not depend on *K*, at the leading order. Moreover, *C*_*EE*_(Δ) and *C*_*II*_ (Δ) depend on the synaptic strengths only because the autocorrelations *A*_*E*_ and *A*_*I*_ depend on these parameters.

Figure 2 depicts simulation results for *N* = 40000 and *K* = 2000. Figure 2a plots *C*_*EE*_(Δ) for 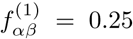 (*α, β E, I*) and two sets of values for the interaction strengths (solid and dashed lines). For comparison we also plots the results of a simulation when the connectivity is unstructured 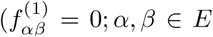 *I*; gray line). In all these cases *C*_*EE*_(Δ) is very small (note the scale on thr y-axis). When the connectivity is spatially modulated *C*_*EE*_(Δ) varies with distance. However, the spatial averages are comparable in the three cases considered 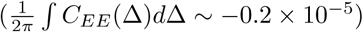. This is in agreement with Eq. (30) since, to the leading order, the autocorrelations, *A*_*E*_ and *A*_*I*_, are not expected to depend on whether the connectivity is spatially modulated or not.

Note that according to Eq. (30), *C*_*EE*_(Δ) and *C*_*II*_ (Δ) are negative for close-by neurons (Δ small) and positive for neurons that are far apart. This is the case for the set of parameters corresponding to the solid line in Fig. 2a (see also Fig. 2b-d). However, when finite *K* corrections are not negligible, which happens when *|T* ^(^1^)^Δ^(^1^)^ *|* is sufficiently small, short range correlations can be positive and longer range correlations negative (Eq. (30)). This is the case in Fig. 2a, dashed line.

**FIG. 2.**
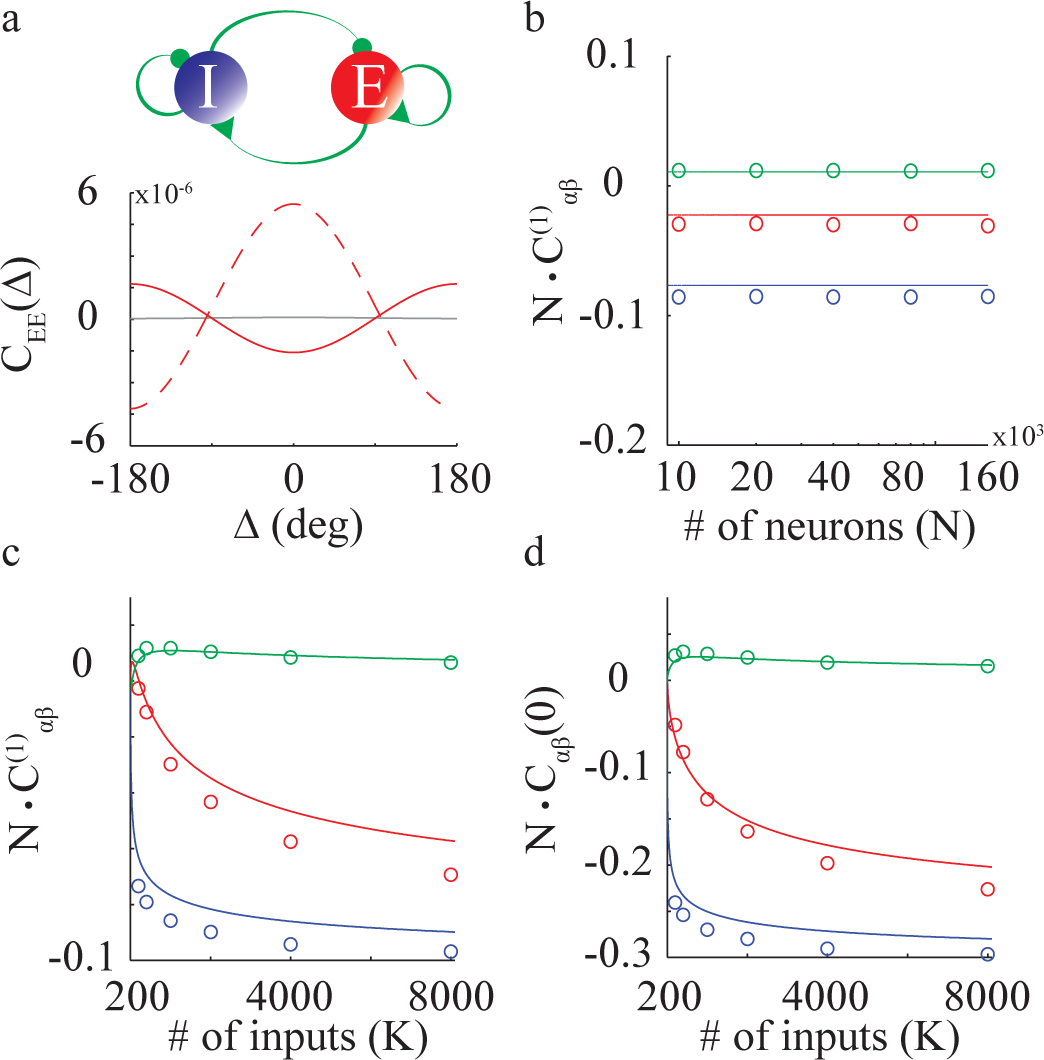
Correlations in two-population E-I networks. Connection probabilities are as in Eq. (29). Panels a, c, d: *N* = 40000. **a.** Top panel: The network architecture. All connections are spatially modulated 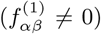. Bottom panel: Simulation results for *C*_*EE*_ (Δ) with *K* = 2000. Red: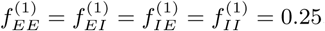. For comparison the correlation is also plotted for 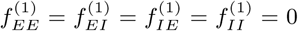 (Gray). For red-solid and gray lines other parameters are *J*_*EE*_ = 0.3, *J*_*IE*_ = 3, *J*_*EI*_ = 2.5, *J*_*II*_ = 5; *I*_*E*_ = 0.3, *I*_*I*_ = 0.3, *T*_*E*_ =1, *T*_*I*_ = 0.7. With these parameters 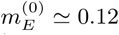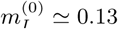 *g*_*E*_ ≃ 0.22, *g*_*I*_ ≃ 0.1 and *A*_*E*_ ≃ 0.1, *A*_*I*_ ≃ 0.1. For red-dashed line other vparameters are: *J*_*EE*_ = 1; *J*_*IE*_ = 2; *J*_*EI*_ = 1; *J*_*II*_ = 1.5; *I*_*E*_ = 0.2; *I*_*I*_ = 0.08. **b. 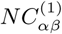** vs. *N* for *K* = 1000. **C.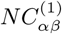** vs. *K*. **d.** *NC*_*αβ*_(0) vs. *K* (*N* = 40000). In panels b, c, d: Solid line: Analytics, Eq. (27). Circles: Simulations. Red: *α* = *β* = *E*. Blue: *α* = *β* = *I*. Green: *α* = *E, β* = *I*. Parameters are as for red-solid line in panel a.

Figure 2b compares simulations (circles) and analytical results (solid lines) for the dependence on *N* of 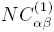 (parameters as in panel a, red solid line). It shows that the spatial modulation of the correlations in the simulation is close to the large *K* analytical results. Figure 2c-d depict the dependence of *NC*_*αβ*_ on *K*. In the whole range of *K* considered here simulations and analytical results are close. *|*For *C*_*αα*_*|* the nearby correlations and modulation amplitudes increase with *K* and are much larger than that of *|C*_*EI*_ *|*.

### IV.2. *T* ^(^1^)^ = Δ^(^1^)^ = 0

We now consider a network in which 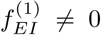 and 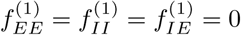. The spatially modulated component of the interaction has therefore an explicit feedforward structure (Fig. 3a, top panel) and *T* ^(^1^)^ = Δ^(^1^)^ = 0.

Solving Eq. (27) shows that the correlations are on average 𝒪 (1*/N*) and that their modulations are

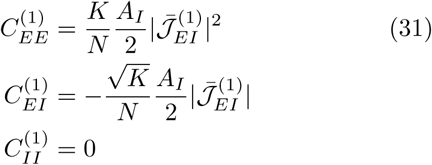

**FIG. 3.**
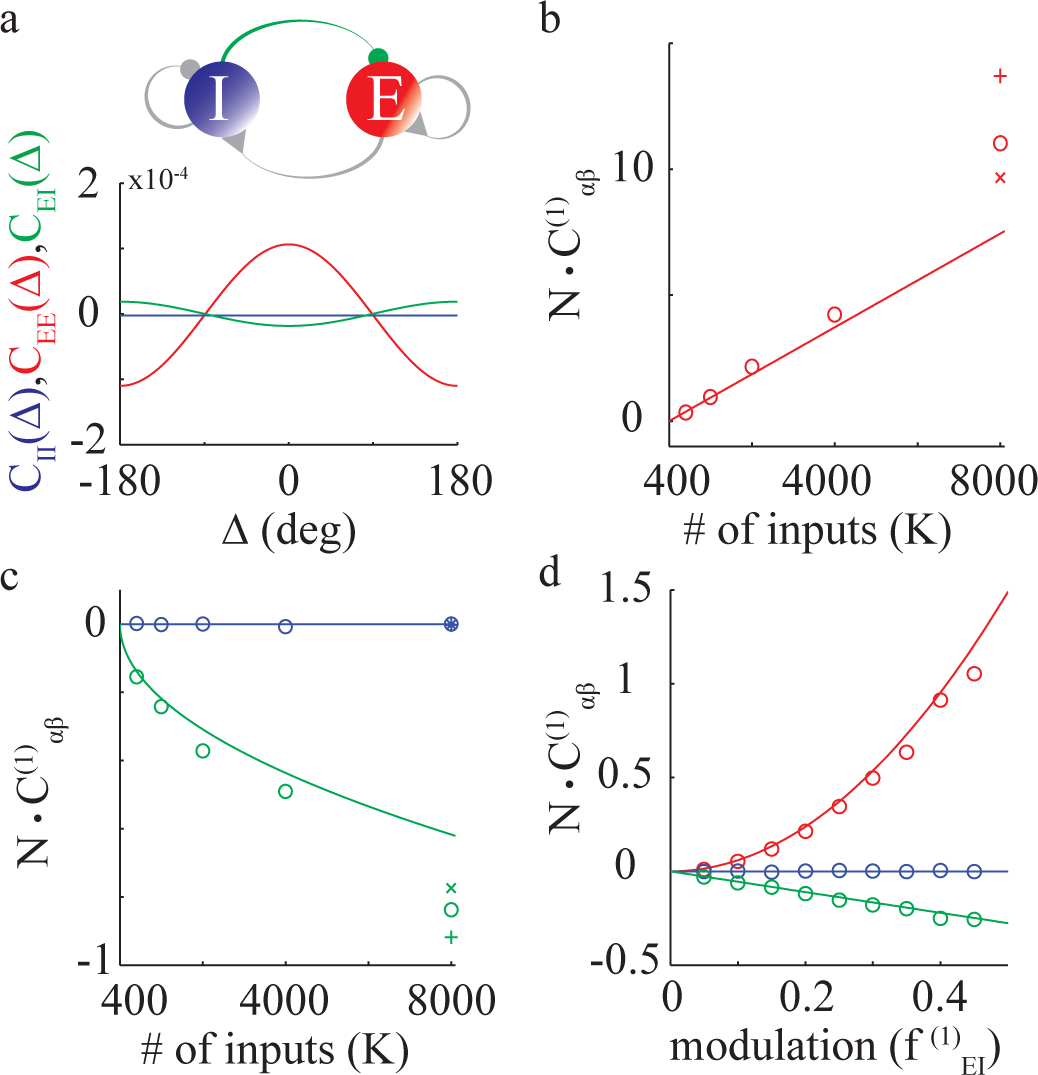
Correlations in a two population network with an explicit feedfworward structure. Connection proba-bilities as in Eq. (29). Parameters: 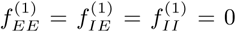 Panels a, b, 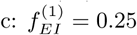. Interaction strengths as in Fig. 2 **a**. Top panel: The network architecture. Green: the connections which are spatially modulated 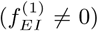. Connections in gray are unstructured 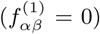. Bottom panel: Simulation results for *C*_*αβ*_(Δ); *N* = 40000, *K* = 2000. **b, c.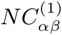** vs. *K*. Solid lines: Analytical results, Eq. (31). Simulation results are plotted for *N* = 20000 (plus), *N* = 40000 (circles) and *N* = 80000 (crosses). **b.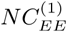**. **c.** Blue; 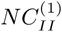. Green: 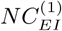 **d.** 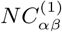 vs.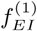. Solid lines: Analytical results, Eq. (31); *N* = 40000, *K* = 400.

As a result, correlations in the E population are spatially modulated and 𝒪 (*K/N*). They are positive for short range and negative for long range. This is in contrast to the correlations in the inhibitory population which are not spatially modulated and 𝒪 (1/N) while the *EI* correlations are spatially modulated and 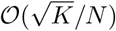.

Figure 3 compares analytical results with simulations. Panel a plots simulation results for 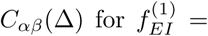 0.25. Other parameters are as in Fig. 2a (grey solid line). Thus, the locally averaged firing rates are the same as in simulations in the latter figure. Correlations in the inhibitory neurons (blue) are extremely weak and are not spatially modulated. In contrast, the correlations of excitatory pairs (red) are larger by two orders of magnitude compared to those in Fig. 2. Correlations between E and I neurons (green) are weaker than those between E neurons. They are negative for nearby neurons while for nearby excitatory pairs they are positive. All these features are in agreement with our analytical results,

The spatial modulation of the EE correlations increases with *K* (Fig. 3b, circles). There is quantitative agreement between simulations and theory (Eq. (31)) up to *K ≈* 4000 for *N* = 40000. In this range,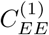 in the simulations varies linearly with *K*. For larger values of *K*,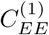 is larger than predicted by Eq. (31). This is because this equation was derived by linearizing the dynamics, which is only valid when correlations are not too large. In fact, simulation results for fixed *K* deviate less from Eq. (31) when *N* is increased (Fig.3b-c). When *K* increases,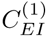, becomes more negative (Fig. 3c; green). Here too, for *N* = 40000 simulations agree well with Eq. (31) up to *K ≈* 4000 and deviations are smaller when *N* is larger. The spatial modulation of the II correlations in the simulations are extremely small (Fig. 3c; blue) as the theory predicts.

Finally, according to Eq. (31), the spatial modulation, 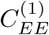, increases quadratically with 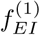 whereas 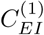 varies linearly with this parameter. Our simulations are in very good agreement with these analytical results (Fig. 3d).

### IV.3. Δ^(^1^)^ = 0, *T* ^(^1^)^ ≠ 0

The network investigated in IV.2 has an explicit feedforward structure. Adding any spatial modulation to the II connectivity (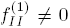, top panel of Fig. 4a) destroys this structure and now *T* ^(^1^)^ ≠0 (while Δ^(^1^)^ ing Eq. (26) for that case, one finds:

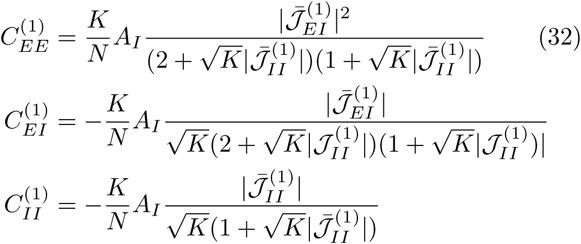

**FIG. 4.**
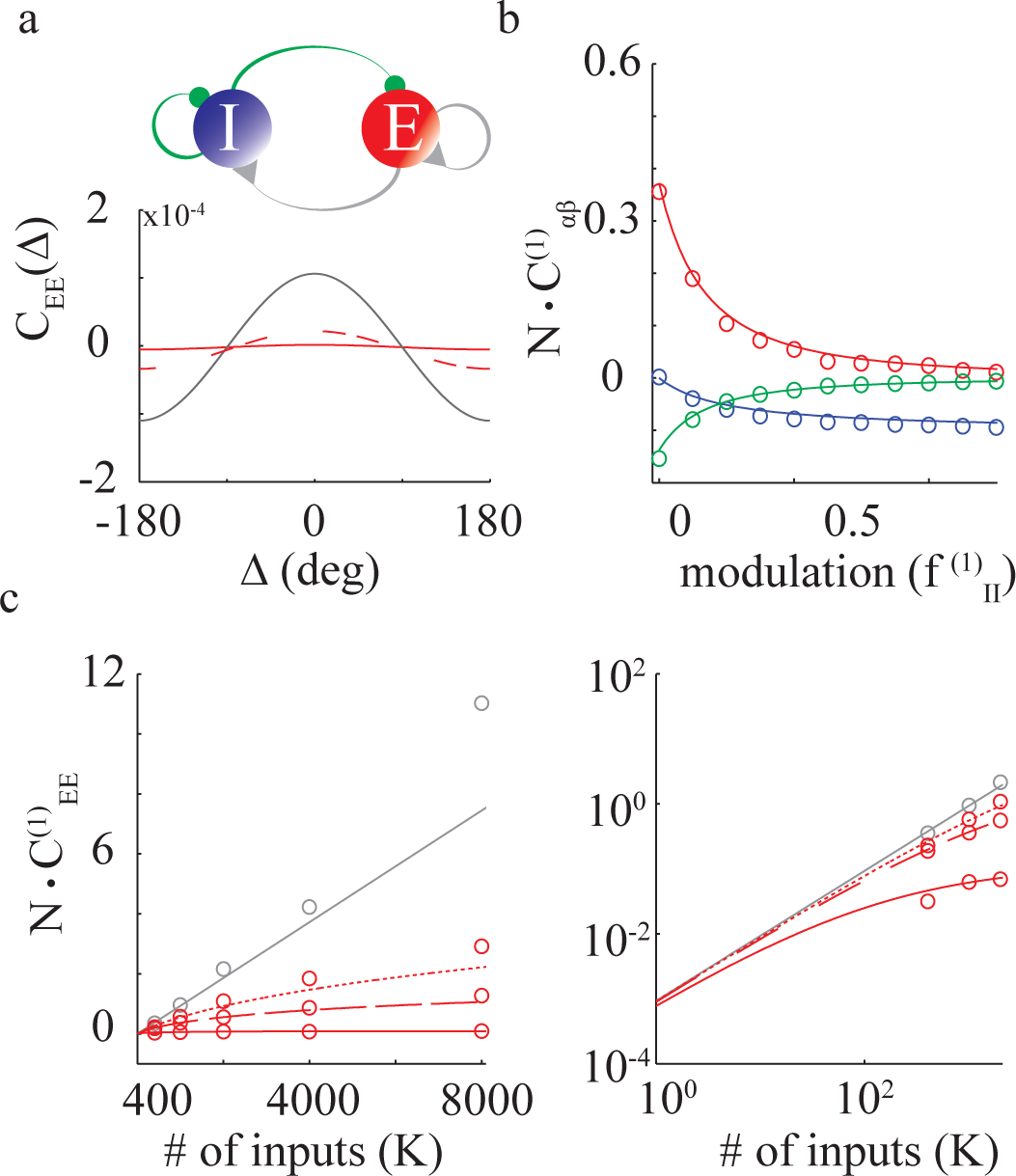
Spatial modulation in the II interactions suppresses the correlations in the E populations. Same network as in Fig. 3 but 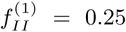. *N* = 40000. **a.** Top panel: The network architecture. Interactions plotted in green are spatially modulated. Bottom panel: Simulation results for *C*_*EE*_ (Δ). *N* = 40000, *K* = 2000. Gray line: 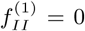; Dashed- red: 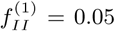; Solid-red: 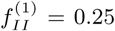. b. 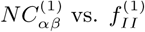. Circles: Simulations. Solid lines: Solution of Eq. (32). *N* = 40000, *K* = 400. Colors are as in Fig. 3. Left panel: 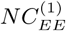 vs. *K*. Solid lines: Eq. (32). Circles: Simulations. Gray line: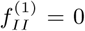; Dotted red: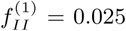; Dashed red: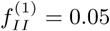; Solid red: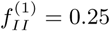. Right panel: Same as left panel but in a log-log scale.

In the large *N, K* limit, *C*_*EE*_(Δ) and *C*_*II*_ (Δ) are both 𝒪 (1*/N*) and do not depend on *K*, whereas *C*_*EI*_ (Δ) is 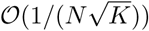. Therefore the addition of a spatial modulation in the II interactions suppresses the correlations that inhibitory projections induce in the E population.

To understand further the origin of this, let us consider the quenched average correlations of the inputs, *Q*_*αβ*_(Δ) (Eq. (19)). Using Eq. (21), one finds

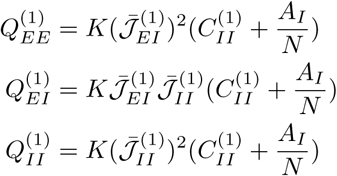

When 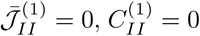, therefore 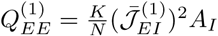. On the other hand, when 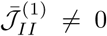, 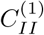 is given by Eq.(32). This yields

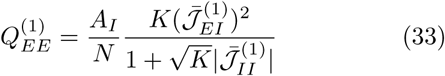

Thus, when II interactions are spatially modulated, a cancellation between terms which are *𝒪* (*K/N*), reduces 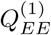 by a factor of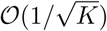. As a result, 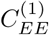 is much smaller than when II interactions are not modulated. A similar argument explains the suppression in 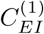.

Fig. 4a depicts simulation results for 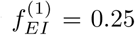 and three values of 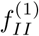. It demonstrates the suppression of correlations in the excitatory population when II interactions are also modulated. The dependence of this effect on 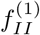 is depicted in Fig. 4b. Increasing 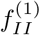, decreases the modulation of all the correlations in very good agreement with the analytical results (compare circles and solid lines).

According to Eq. (32), the correlation in the E population always increases linearly with *K*, for small *K*. The cross-over between this regime, where *C*_*EE*_(Δ) is *𝒪* (*K/N*), and the large *K* regime, where *C*_*EE*_(Δ) is *𝒪* (1*/N*), occurs for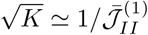. Figure 4c depicts this crossover in numerical simulations. Thus, although in the large *N, K* limit a transition from *𝒪* (*K/N*) to *𝒪* (1*/N*) correlations occurs as soon as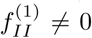, this qualitative difference is significant only when *K* is sufficiently large. In other words, for moderately large number of inputs per neuron, correlations can exhibit a close to linear increase even if the structure of spatial modulation of the interaction matrix is not completely feedforward.

## V. SCALING CORRELATION THEOREMS

In the previous section we studied networks with two neuronal populations. In this case, it is straightforward to analytically derive explicit expressions for the correlations. These expressions are simple enough to fully classify how the network structure affects the scaling (with *K* and *N*) of the correlations. For networks consisting of more than two populations, analytical expressions for the correlations can in principle be derived. However, dealing with these expressions becomes rapidly impractical as the number of populations increases. In the following we adopt an alternative approach. We prove two general theorems, which, given the network architecture, allow us to determine how correlations vary with *K*, without computing these correlations explicitly.

To prove these theorems, we rewrite Eq. (26) in the Jordan basis of 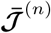. We write

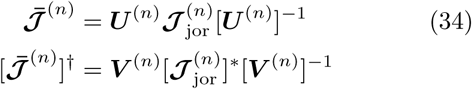

Here *x*^*\**^ denotes the complex conjugate of *x*, ***U*** ^(n)^ (***V*** ^(n)^) are matrices whose rows are the generalized eigen-vectors of 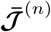 and 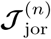 is the Jordan normal form of 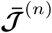. This implies that 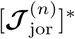 is the Jordan form of where we have defined

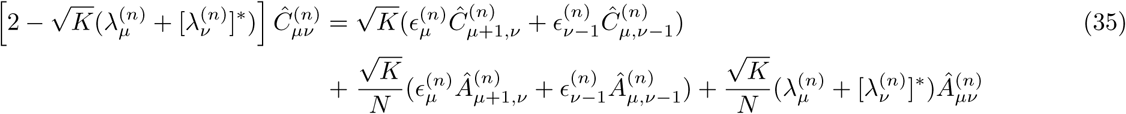

Note that while the matrix ***C***^(n)^is symmetric and the matrix ***A***^(n)^is diagonal, this is not in general the case for *Ĉ*^(n)^and Â^(n)^.

Let us assume that the network is in a stable balanced state in which the matrix Â^(n)^. has no zero elements. In Appendix B we prove

*Correlation Theorem 1:* The *n*^*th*^ Fourier mode of the correlation matrix scales as *𝒪* (1*/N*) if and only 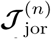 does not have a Jordan block whose real part is a shift matrix (A shift matrix, ***S***, of dimension *P* is a *P × P* matrix of the form *S*_*μv*_ = *δ*_*μ*+1,*v*_).

*Correlation Theorem 2:* If 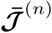 has at least one Jordan block whose real part is a shift matrix, the *n*^*th*^ Fourier mode of the correlation matrix is *𝒪* (*K*^*P*^ ^(n)*-*^1^^*/N*), where *P* (n)is the dimension of the largest block in 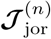, whose real part is a shift matrix.

*Corollary:* The *n*^*th*^ Fourier mode of the correlation matrix is *𝒪* (*K* ^*D-*^1^^*/N*) if and only if 𝒯^(n)^ is nilpotent of degree *D*.

In Appendix B we also show how to extend these re-sukts to case where Â^(n)^. has zero elements.

The matrix ***U*** ^(n)^can be viewed as a transformation of the original network of *D* populations into a network of *D* effective populations. The condition that 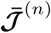 has a Jordan block which is a shift matrix of size *P*, can be interpreted as the existence of *P* effective populations whose effective interaction matrix is feedforward in its *n*^*th*^ Fourier mode. In other words, the original network exhibits a hidden feedforward structure, which is embedded in the *n*^*th*^ Fourier mode of its connectivity. Theorems 1 and 2 therefore implies that only when such a structure exists, the *n*^*th*^ mode of the correlation matrix increases with *K*. To know which elements in this matrix increase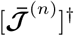 is the 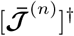 We can write

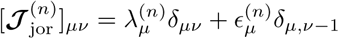

with *μ, v ∈* 1, *…, D* and 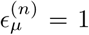 within a Jordan block and zero otherwise.

Equation (26) then yields

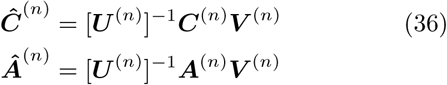

with *K* one has to compute the matrices ***U*** ^(n)^and ***V*** ^(n)^ (see Eq. (36)).

## VI. APPLICATIONS OF THE CORRELATION THEOREMS

In this section we consider networks comprising *D* populations with connection probabilities

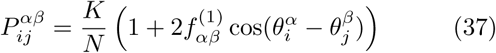

with *α* = 1, *…, D* and *β* = 1, *…, D*.

### VI.1. Two population networks

For a network of two populations the Jordan form of the matrix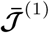 has the form: 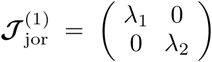if 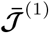 is diagonalizable. Otherwise, it has the form: 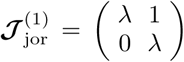 with *λ* real. Theorem 1 and 2 imply that only in the second case with *λ* = 0, some of the correlations *C*_*αβ*_(Δ) are *𝒪* (*K/N*). Otherwise, all correlations are *𝒪* (1*/N*). It is equivalent to say that some correlations are *𝒪* (*K/N*) if and only if 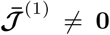and *T* ^(^1^)^ = Δ^(^1^)^ = 0. We therefore recover the result we derived in Section IV without explicit computation of the correlations.

As noted above, the correlation theorems do not tell us which of the elements of the correlation matrix are *𝒪* (*K/N*) when 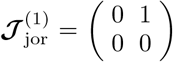. However, since a 2 *×* 2 matrix has such a Jordan form if and only if it is nilpotent **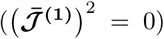** we have to consider two types of connectivity:

Type 1: The network has an explicit feedforward structure, *i.e.*, the interaction matrix is either 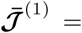 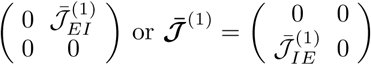.

In the former case, the matrix ***U*** ^(^1^)^ is 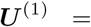 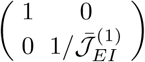, whereas 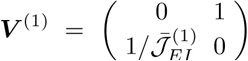. Then Eq. (36) gives (see Appendix B, Eq. (B8))

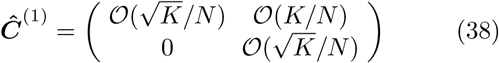

Using Eq. (36), one finds that 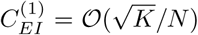, 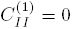, whereas 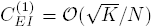, in agreement with Eq. (31). A similar calculation in the latter case (when the modulation is in the IE interactions) gives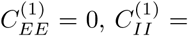, 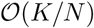, and 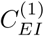, is 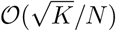 all in agreement with Eq. (27).

Type 2: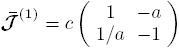 where *a, c >* 0.

In this case the network has no *explicit* feedforward structure since all four interactions (*EE, EI, IE, II*) are spatially modulated. It has, however, a *hidden* feedforward structure, as revealed by the Jordan form of the interaction matrix.

For this network, the transformation matrices are 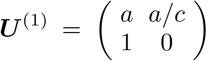 and 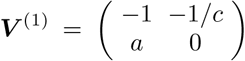. Using Eq. (36) and Eq. (38), it is clear that the transformation from *Ĉ*^(^1^)^ to C^(^1^)^ mixes elements which are 0 and 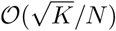, with those which are *𝒪* (*K/N*). Thus, while in 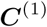only the element 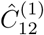is *𝒪* (*K/N*), all the elements of the correlation matrix C^(^1^)^ are *𝒪* (*K/N*). This is also in line with Eq. (27).

We consider an example of such a network in Fig. 5. The parameters *J*_*αβ*_ and the external inputs are as in Fig. 2-4. Therefore, to leading order, the population averaged activities, *m*_*α*_, the autocorrelations, *A*_*α*_, and the population gains, *g*_*α*_ are the same as in Figs. 2-4. The modulation of the connection probability, 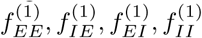, are all non-zero (Fig. 5a) and tuned so that:

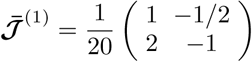

**FIG. 5.**
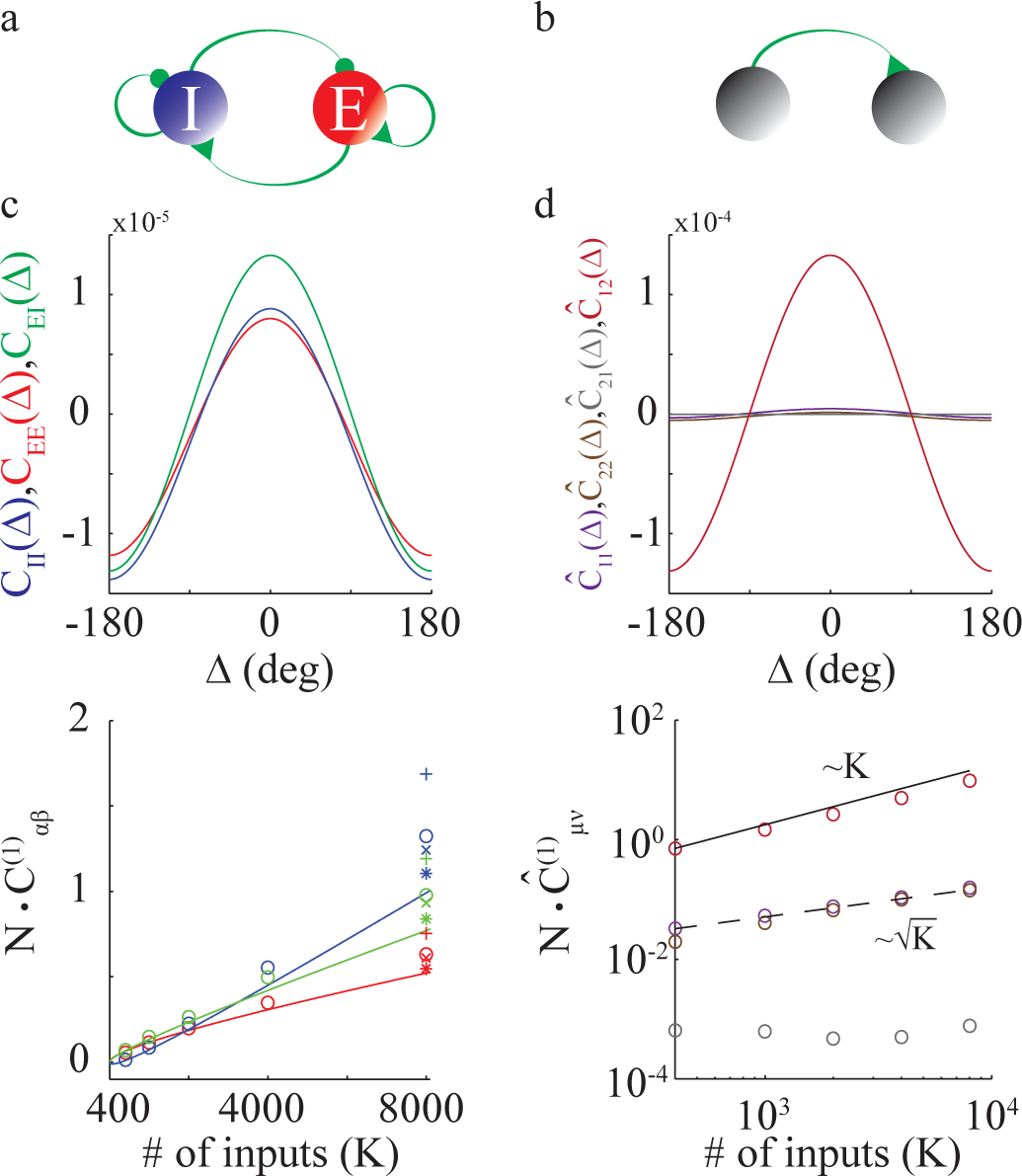
Correlations in a two population network with a hidden feedforward structure. **a.** Network architecture. All connections are modulated. Connection probabilities are as in Eq. (29) with 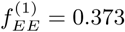 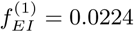 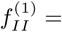 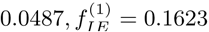. Connection strengths are as in Figs. 2-3. **b.** Hidden feedforward structure of the mode *n* = 1 as revealed in the Jordan basis. **c.** Top panel: Simulation results for *C*_*αβ*_(Δ). *N* = 40000, *K* = 2000. Bottom panel: 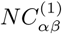 vs. *K*. Solid lines: Analytical results (Eq. (27)). Simulations are plotted for *N* = 20000 (plus), *N* = 40000 (circles), *N* = 80000 (cross) and *N* = 160000 (asterisks). **d.** Top panel: Correlation matrix in the Jordan basis of 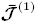. Purple: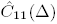; Brown: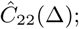; Gray: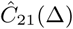; Red: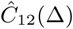. Bottom panel: 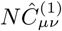 vs. *K* in log-log scale. Solid line: Linear fit. Dashed line: Fit to a square root function.

The Jordan form of this matrix is graphically represented in Fig. 5b. The top panel in Fig. 5c depicts simulation results for the correlations in this network. They are all positive for close-by neurons and their spatial modulations increase approximately linearly with *K* (Fig. 5c, bottom). The top panel of Fig. 5d shows that most of the power in these correlations results from the element 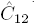 (red). The latter increases l<u>in</u>early with *K* (Fig. 5d, bottom), while 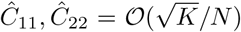 and *Ĉ*_21_ is two order of magnitude smaller and does not exhibit significant change with *K* (note the log-log scale in this figure). These simulations are in quantitative agreement with Eq. (27) up to *K ≈* 4000 and the deviations for larger *K* decrease with *N* (as in Fig 4).

The interaction matrix, 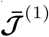, depends on the matrix ***J***^(^1^)^, and on the population gains, *g*_*E*_ and *g*_*I*_. If the connectivity matrix has an explicit feedforward structure, the interaction matrix has also such a structure, independantly of the population gains. Thus, although changing the external inputs, *I*_*E*_, *I*_*I*_, modifies this gains, this does not destroy the feedforward structure and thus does not change the scaling of the correlations with *K* and *N*. In contrast, in networks with a hidden feedforward structure, this scaling is sensitive to perturbations in the external inputs since the hidden feedforward structure is destroyed unless the ratio *g*_*E*_*/g*_*I*_ remains constant.

### VI.2. Examples with three populations or more

The two networks depicted in Fig. 6 consist of one excitatory and two inhibitory populations. The positivity of the spatial averaged activity of these three populations, 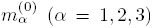, constrains the parameters (see Section III), *J*_*αβ*_, through a set of inequalities. We leave the calculation of these conditions to the reader.

In both cases, the first Fourier mode of the population average connectivity matrix has the form

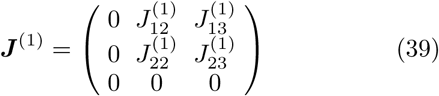

**FIG. 6.**
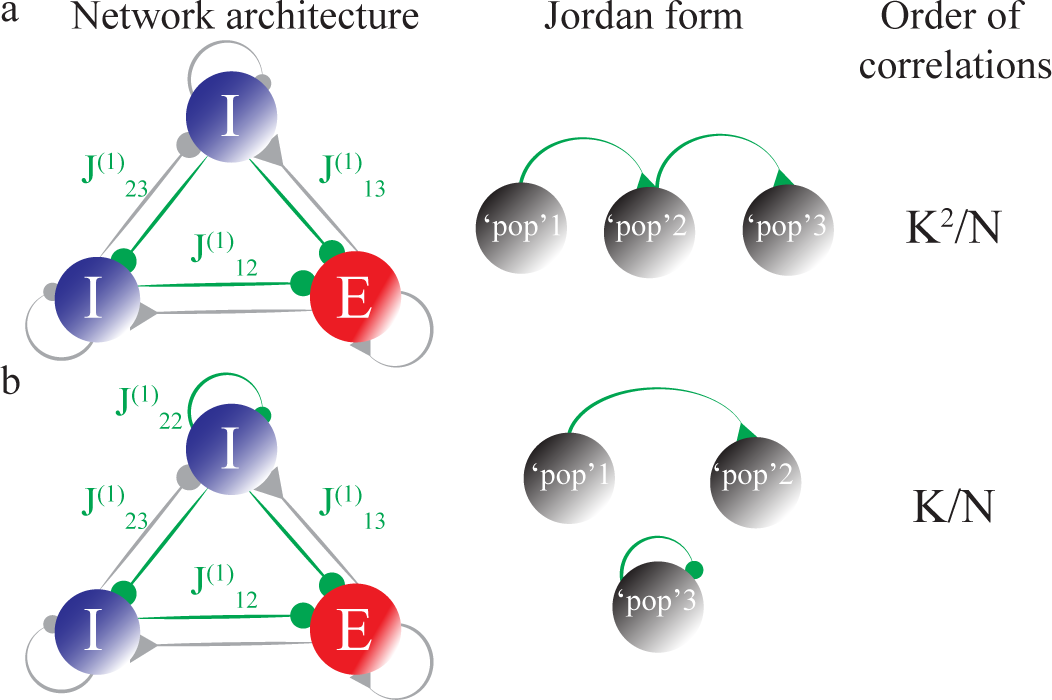
Examples of networks with three populations and their Jordan representations. Probability of connections are as in Eq. (37). **a-b.** Left: A network of three populations, two inhibitory and one excitatory. Gray: Unstructured connections. Green: Spatially modulated connections. Middle: The Jordan representation of the *n* = 1 Fourier mode of the network connectivity (left panel). Right: The scaling of the strongest correlations. **b.**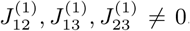. Other entries of the matrix ***J***^(^1^)^ are zero (see main text). **b.** Same as (a), but with 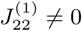.

with 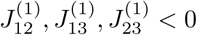 (left panels of Fig 6a-c).

In the network of Fig. 6a,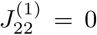. Therefore ***J***^(^1^)^ and 𝒯^(^1^)^ are nilpotent of degree 3. According to the Corollary of Section V, the first mode of the correlation matrix is *𝒪* (*K*^2^*/N*).

The Jordan form of 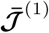 is

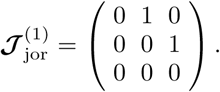

This is graphically represented in Fig. 6a, middle panel. The matrix *Ĉ*^(^1^)^ satisfies (see Appendix B, Eq.(B8))

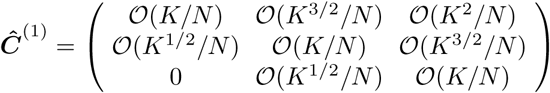

Using the transformation matrices (Eq.(34)), one can show that correlations are *𝒪* (*K*^2^*/N*) only within the excitatory population.

In the network in Fig. 6b,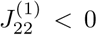. Therefore, the interaction matrix, 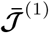, is not nilpotent. Its Jordan form is

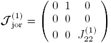

The upper Jordan block is a shift matrix of degree 2. The corresponding feedforward structure is graphically represented in Fig. 6b (middle panel). According to The orem 2, *Ĉ*^(^1^)^ is 𝒪 (*K/N*). It satisfies (see Appendix B, Eq. (B8))

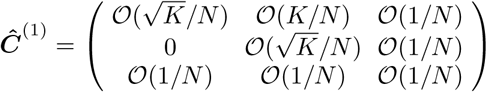

and using the transformation matrices one can show that only 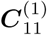 is *𝒪* (*K/N*). Other correlations are either 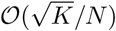 or *𝒪* (1*/N*).

This approach can be generalized to arbitrary number of populations to classify the scaling of the correlation matrix for different architectures. Examples of networks with four populations are depicted in Fig 7, together with the graphic representations of their Jordan forms and the maximum order of the correlations.

## VII. CONSTRAINT ON SCALING OF THE NUMBER OF INPUTS WITH THE NETWORK SIZE

In this section we assume that *K* and *N* scale together

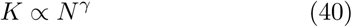

with 0 *< γ ≤* 1.

Equation(26), which determines the correlations of the neuronal activities, is obtained under the Ansatz that the correlations between the inputs 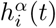 and 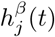 are sufficiently small, namely *o*(1) (see Appendix A). This condition is more stringent than the balance correlation equation, Eq. (23). It can be written as

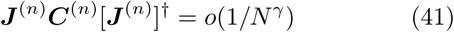

This condition constrains *γ* as we now show.

According to Theorem 1, if the Jordan form, 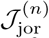, has no Jordan block whose real part is a shift matrix for any *n*, correlations will be *𝒪* (1*/N*). This will also be the order of ***J***^(n)^***C***^(n)^[***J***^(n)^]^*†*^. Therefore, Eq. (41) only requires *γ < γ*_max_ = 1.

If the Jordan form,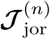, contains a Jordan whose real part in the shift matrix, we have to apply Theorem 2. In this theorem, *P* (n)is the dimension of the largest block in 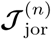, with a real part which is a shift matrix (see Section V). Let us denote by *P*_max_ the largest *P* (n)over all Fourier modes, i.e.,

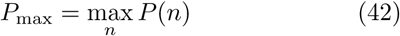

Equation (41) implies

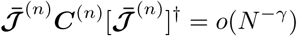

for all *n*. This yields in the Jordan basis

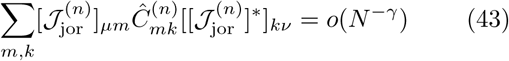

for all *μ, v*.

By definition of *P*_max_, for at least one Fourier mode, *n*, the matrix 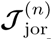 has at least one block whose real part is a shift matrix of degree *P*_max_. In general, there can be several such Jordan blocks. For example, in the network depicted in Fig. 7b, for which *P*_max_ = 2, there are two Jordan blocks with *P* = 2.

**FIG. 7.**
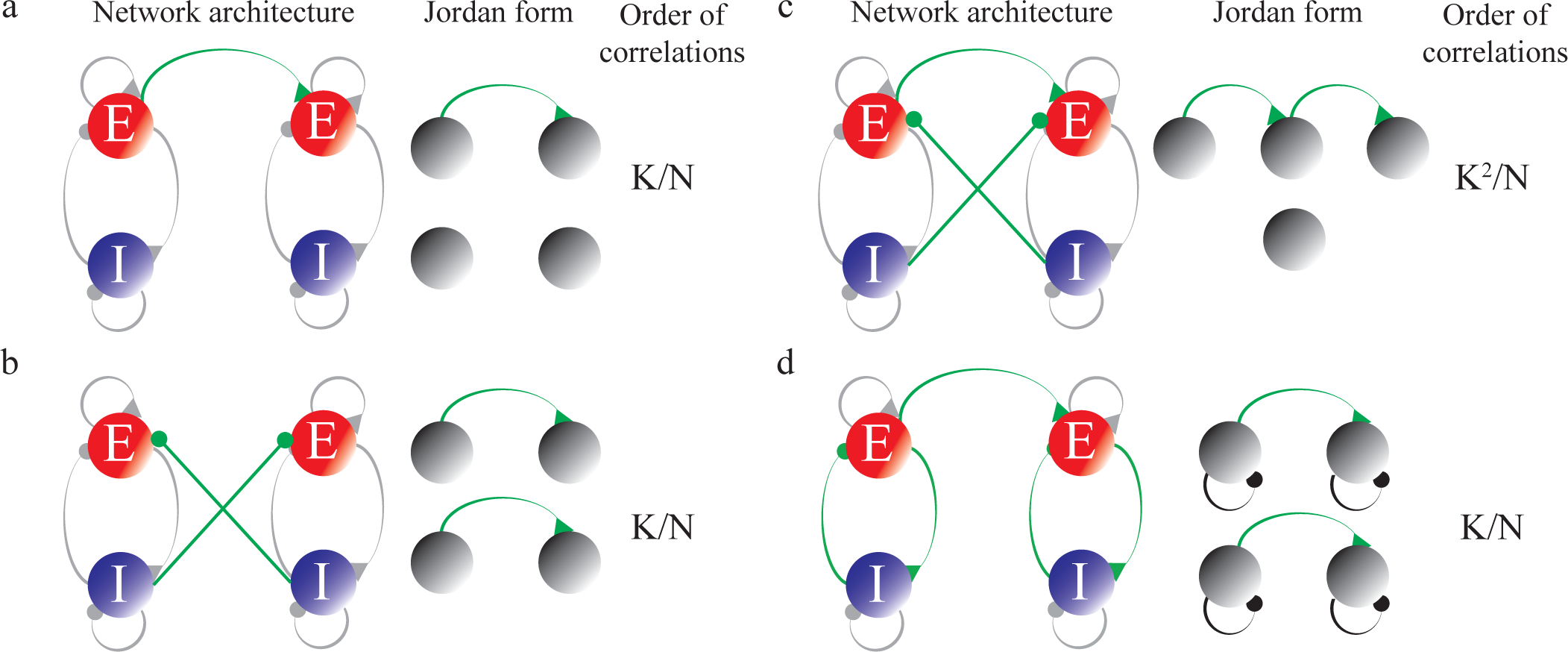
Examples of networks with four populations and their Jordan representations. Probability of connections are as in Eq. (37). **a-c.** Left: The network consists of two coupled E-I networks. Gray: Unstructured connections. Green: Spatially modulated connections. Middle: The Jordan representation of the *n* = 1 Fourier mode of the connectivity (network of the left panel. Right: The scaling of the strongest correlation. **d.** Same as (a-c), with a population averaged connectivity matrix as in Eq. (46). Middle: The Jordan form is complex (Eq. (47)). Black lines corresponds to complex eigenvalues.

We first assume that all blocks which are a shift matrix of size Pmax are real. Since for such blocks in 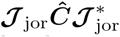 scale the same with K, it is sufficient to consider the case where there is only one such block. We denote it by S and by *Ĉ*_max_the corresponding block in *Ĉ*. Equation (41) then yields

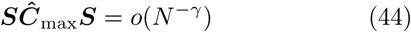

For instance, for *P*_max_ = 2, we have (see Eq. (B8))

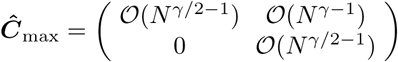

and thus

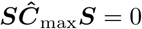

Therefore, for this block Eq. (41) is always satisfied. The latter equation, however, also applies to other Jordan blocks and Fourier modes. This implies that *γ < γ* _max_ =1

For *P*_max_ = 3, we have

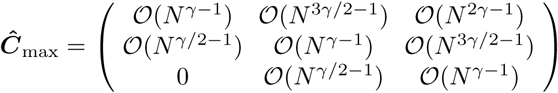

and

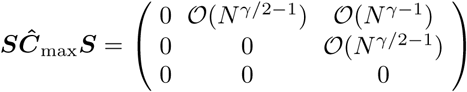

Thus, Eq. (41) is satisfied only if *γ < γ*_max_ = 1*/*2.

In general, for a *P* _max_ *× P* _max_ shift matrix *S****Ĉ*_max_*S*** is 𝒪 (*N* ^*γ*(*P* max*-*^2^)*-*^1^^). This implies that *γ < γ*_max_ with

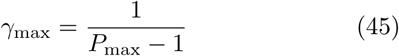

Let us now consider networks in which there is at least one pair of complex conjugate Jordan blocks whose real parts are a shift matrix of size *P*_max_. An example, of such a network is depicted in Fig. 7d. The first Fourier mode of the population averaged connectivity matrix in this example is

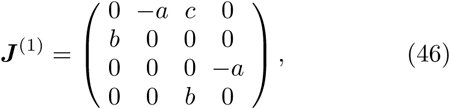

with *a, b, c* real and positive. For this network

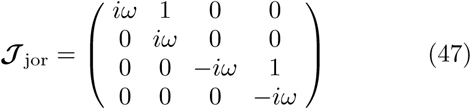

with 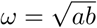.

These complex conjugate blocks can in general be written as ±*iv****I*** + ***S***, where ***I*** is the identity matrix of size *P*_max_. Thus

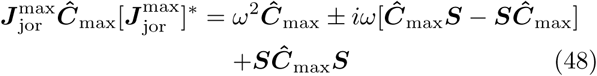

As shown above, *Ĉ*_max_= *𝒪(N* ^*γ*(P^_max_^-^1^)-^1^^)and S *Ĉ*_max_ *S=𝒪(N* ^*γ*(P^_max_^-^2^)-^1^^). It is straightforward to also show that *Ĉ*_max_*S–SĈ*_max_*=𝒪(N* ^*γ*(P^_max_^-^3^/^2^)-^1^^). Therefore 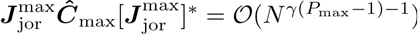. Equation (41) is then satisfied only if *γ < γ*_max_, with

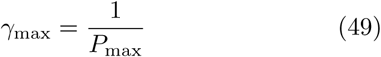

According to Theorem 2, if ^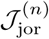^ has a block whose real part is a shift matrix for at least one mode *n*, ***C*** = 𝒪 (*N* ^*-α*^) where *α* = 1 − *γ*(*P*_max_ *-* 1). If *γ < γ*_max_, correlations in the activity will decrease more slowly than 1*/N*, when *N* is increased. If *γ > γ*_max_ correlations will increase with *N* and the network will not operate in the balanced regime. Finally, if *γ* = *γ*_max_, our theory will give (1) correlations in the input which is inconsistent with the Ansatz in Eq. (41). In this case substantial corrections to the Eq. (26) should be taken into account. A different approach, similar to the one in [28] must be adopted to self-consistently determine these correlations.

## VIII. DISCUSSION

### VIII.1. Main results

We developed a theory for the emergence of correlations in strongly recurrent networks of binary neurons. Each neuron receives on average *K* inputs from each of *D* populations, with probabilities which are spa<u>tia</u>lly modulated. The synaptic strengths scale as 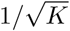 and the network operates in the balanced regime [45]. For simplicity we considered networks with a one dimensional ring architecture with connection probabilities solely dependent on distance and on the nature (excitatory or inhibitory) of the preand postsynaptic populations.

We present a balanced correlation equation, which together with the balanced rate equation, define the balanced regime and insure that mean inputs to the neurons and their fluctuations are both 𝒪 (1). We derive a set of equations that determine the equilibrium values of the quenched averaged correlations and we show that the solution of these equations is stable provided that the solution of the balanced rate equations is stable.

Key results of our work are two scaling correlation theorems. The first shows that generically, all the Fourier modes of the quenched average correlations are small when *N* and *K* are large. They are of 𝒪 (1*/N*), and independent of *K* to leading order. This is true in the large *N* limit even if we take *K* = *pN*, provided that *p* is not too large. However, the second theorem states that there are recurrent network architectures in which some of the Fourier modes of the quenched averaged correlations increase with *K*. These architectures are characterized by an explicit, or a hidden, feedforward structure in those modes. This structure is revealed by the Jordan form of the interaction matrix averaged over realizations. If this Jordan form contains a block whose real part is a shift matrix of size *P >* 1, the corresponding mode in the correlation increases at least as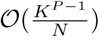. Importantly, in these cases the network still operates in the balanced regime provided that 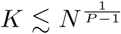 (or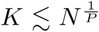, see Section VII). Corrections to the theory become important when *K* approaches 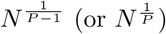.

### VIII.2. Generality of the results

For simplicity, we assumed that synaptic weights depend only on the identities of the populations to which pre and postsynaptic neurons belong. The results, however, will not change if the weights are heterogeneous with distributions that depend on the pre and postynaptic populations, as long as the mean and the variance of all these distributions are finite.

For notational simplicity we assumed that all populations have the same number of neurons, *N*, and each population receives, on average, inputs from *K* neurons in every population. The theory can be easily extended to networks in which population *α* has *N*_*α*_ = *v*_*α*_*N* neurons and the average number of connections from population *β* to population *α* is *K*_*αβ*_ = *κ*_*αβ*_*K*. This will not affect the scaling of the correlations with *N* and *K*. Prefactors, however, will be different. For instance, assuming four times fewer inhibitory than excitatory neurons in the two-population networks of subsection IV.1, without changing the number of connections per neuron, the correlations of the inhibitory neurons will increase by a factor of 4.

We focused on networks with a one-dimensional ring architecture and connection probabilities which are solely distance dependent. This greatly simplifies the problem, because when averaged over realizations, correlations depend solely on distance. Furthermore, because of the linearity of the self-consistent equation for the correlations, the different Fourier modes decouple, allowing us to analyze each mode separately. However, our analytical approach does not require rotation invariance. It can be extended to any network architecture for which the Jordan normal form of the interaction matrix, averaged over realizations, can be established.

### VIII.3. Robustness and self-consistency of the results

The theory presented here makes the Anzatz that correlations are sufficiently small so that the dynamics of the crosscorrelations can be linearized (Appendix A). If this Anzatz is correct, the theory is self-consistent. When the theory predicts correlations which are 𝒪 (1), non-linear terms contribute and the correlations start to deviate from the theoretical value (see for example simulation results for *N* = 40000 in Fig. 3b). Nevertheless, the order of the correlations is still correctly predicted.

In the large *N, K* limit, only networks with feedforward structures -hidden or explicit- can exhibit correlations that increase with the average number of inputs. However, when *K* and *N* are only moderately large, as is the case in biological systems, a strict tuning of the architecture is not necessary. This is because there is a crossover between the regimes of strong and weak correlations as *K* is increased (see Fig. 4c). As shown in Appendix B1, the value of *K* for which this crossover occurs depends on the eigenvalues of the interaction matrix.

### VIII.4. Relation to previous works

Non-interacting neurons can exhibit correlations if they share feedforward inputs [57, 58]. For instance, Litvak et al. [59] investigated a chain of layers of integrateand-fire neurons lacking any recurrent interactions and coupled only feedforwardly. In their model, each neuron in a layer receives inputs from the same number of excitatory and inhibitory neurons in the previous layer in such a way that their temporal averages *exactly* balance. They found a build up of correlations along the chain. This is because the correlations induced by shared feedforward inputs are not suppressed during the activity propagation since the network lacks any recurrent interactions and is thus purely feedforward.

Cortes and van Vreeswijk [60] studied a chain of strongly recurrent unstructured E-I subnetworks coupled through excitatory unstructured feedforward projections. They found a gradual build up of correlations along the chain. These correlations, however, decrease if the connectivity, *K*, and the sub-networks size, *N*, increase together. This result is in agreement with our theory which predicts that the correlations are 𝒪 (1*/N*) through the whole chain.

In [43] Ginzburg and Sompolinsky considered networks of binary neurons with finite temperature Glauber dynamics, unstructured, dense (*K* = *pN*) connectivity and weak i<u>nte</u>ractions, i.e., of the order of *𝒪* (1*/K*) and not 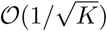 as in our work. Mean Field theory shows that in these networks correlations are (1*/N*) with a prefactor which diverges in the zero temperature limit. This is in contrast to what happens in the strongly recurrent unstructured networks we considered here where correlations also scale as 𝒪 (1*/N*) but with a prefactor, which is finite despite the fact that we assumed zero temperature Glauber dynamics. This is because in strongly recurrent networks, intrinsic noise emerges from the deterministic dynamics of the network.

They also demonstrated that in their model the correlations amplify up to 𝒪 (1) at Hopf bifurcation onsets [43]. At such onsets the dynamics exhibit critical slowing down and thus this amplification is accompanied by a divergence of the decorrelation times. Our work demonstrates a different amplification mechanism: it occurs at a point where the Jordan form of the interaction matrix, 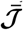 contains a block which is a shift matrix. Since there is no critical slowing down at such a point, the decorrelation times are finite and on the order of the update time constant (data not shown). Our theory may thus account for substantial correlations with short time scales, as frequently observed in the brain [28, 31, 61].

Renart et al. [28] and Helias et al. [49] investigated strongly recurrent unstructured networks with one excitatory and one inhibitory population of binary neurons which are densely connected, i.e with *K ∝ N*. They found that in these networks mean pair correlations were 𝒪 (1*/N*). As they only considered dense connectivity they could not, however, disentangle the dependence on *N* and *K*. Here, we considere different relations between *K* and *N* and show that in unstructured networks the correlations are 𝒪 (1*/N*), and in practice do not depend on *K*. This last result is remarkable since one would expect correlations to increase with the degree of connectivity. This is not the case: the balance of excitation and inhibition prevents that to occur in unstructured networks.

On the other hand, and somewhat surprisingly, we found that even if the fraction of common inputs shared by neurons is very small, a build up of correlations can still occur for some network architectures. For example, in a network of four populations with a feedforward structure, correlations would be of 𝒪 (*K*^3^*/N*), and thus, to satisfy the balanced correlation equation, the scaling of *K* with the network size can be at most *K* = 𝒪 (*N* ^1^*/*^3^). However, with this architecture and scaling, the probability of two neurons to share their inputs is 𝒪 (*N* ^*-*^4^*/*^3^^) (Eq. (17)) and the number of inputs shared by two neurons is therefore 𝒪 (*N* ^*-*^1^*/*^3^^). Thus, although the number of shared inputs goes to zero in the large N limit, correlations get amplified up to (1) thanks to the feedforward architecture.

Rosenbaum et al. [39] have recently investigated how feedforward excitation can drive correlations in spatially structured E-I networks operating in the balanced regime. The specific architecture they considered is reminiscent of the particular example presented in Fig. 7a. In their study the fluctuations which drove the correlated activity were those in the feedforward inputs and the contribution to the correlations of the fluctuations generated by the recurrent dynamics was neglected. In contrast, our work focuses on the role of the *recurrent* dynamics in the emergence of correlations.

We recently studied the emergence of correlations in a network consisting of two strongly recurrent E-I subnetworks, the first projecting to the second with topographically organized feedforward connections [17]. The architecture in that work is also reminiscent of the example presented in Fig. 7a. We showed that while in the first subnetwork correlations were weak, 𝒪 (1*/N*), in the second subnetwork the activity was self-organized in such a way that correlations in macroscopic sub-populations of excitatory neurons were finite and did not depend on *N* and *K* when the latter were sufficiently large. This does not contradict our theory. In the architecture considered in [17] subsets of neurons in the first subnetwork are projecting to a macroscopic fraction of neurons in the second subnetwork. This organization generates correlations between elements of the connectivity matrix, unlike in the models considered here. Generalizing our theory to such cases is possible but beyond the scope of this paper.

### VIII.5. Directions for future work

The theorems of Section V, tell us how the Fourier modes of the averaged correlations scale for large *N* and *K*. In the examples considered in Sections IV, VI and VII, we focused on connectivities whose Fourier expansion involve only two modes. In these cases, the scaling of the spatial correlations can be immediately deduced from that of the Fourier modes. This will also be the case for interactions described by a finite number of Fourier modes. However, if the interactions are described by an infinite number of modes, inferring the scaling of the correlations from those of their modes can be more complicated. For instance, if one takes the large *N* limit with *K ∝ N*^*γ*^, the convergence of the Fourier series may not be uniform in *N*. In that case the scaling with *N* of the spatial correlations may be highly non-trivial. We will address this issue in an upcoming paper.

The present paper focuses on locally averaged twopoint correlation functions. Recent progress in experimental techniques will create large data sets of neuronal activities from which distributions of pairwise correlations can be extracted. The power of theoretical approaches to interpret such data will be greatly enhanced if they provide not only locally averaged correlations but also higher order statistics of their distributions. Thus it would be interesting to extend our approach to estimate the scaling of higher order moments of correlations in binary networks.

Several previous studies investigated EI networks (*e.g*. [49, 62]) in which inhibition and excitation were unstructured and their strengths were only a function of the presynaptic neurons, i.e., *J*_*EE*_ = *J*_*IE*_ and *J*_*II*_ = *J*_*EI*_. Other studies assumed unstructured connectivity with *J*_*αE*_ = *-J*_*αI*_, *α ∈ {E, I}* (*e.g.* [63]). In both cases the interaction parameters are on the edge of the region where the network evolves towards the balanced state (see Eq. (11)). The network dynamics may be qualitatively different on the edge of this region than inside it. It is thus not clear that the scaling theorems presented here apply to these tuned cases. A further investigation of the correlation structure in such networks is a subject for future research.

Do the conclusions derived here for networks of binary neurons hold for networks with more realistic single neuron dynamicsv To approach this question we performed extensive numerical simulations of strongly recurrent networks consisting of one inhibitory and one excitatory population of leaky integrate-and-fire (LIF) neurons. The detailed analysis of these simulations will be presented elsewhere. In brief, we found in our simulations, that in these networks the averaged pairwise correlations scale with *K* and *N* in manner that is consistent with the theory presented here for binary networks. It would be very interesting to extend our analytical approach to these type of networks.

To conclude, van Vreeswijk and Sompolinsky [45, 46] investigated strongly recurrent networks of binary neurons with unstructured sparse connectivity. They showed how these network dynamics evolve into a balanced state. Due to the sparseness of the connectivity, correlations are negligible in these networks. Subsequent studies [28, 49] extended these results to unstructured networks with dense connectivity, and found that here too the network dynamics evolve to a balance state in which reverberations keep the correlations very small. In contrast, as shown here, in networks with structured connectivity correlations can be large. The theory presented here gives the conditions on the network architecture to evolve into a balanced state with strong correlations and show how they depend on the network size and number of connections.

## ACKNOWLEDGMENTS

We thank Gianluigi Mongillo and German Mato for fruitful discussions. Work conducted in the framework of the France Israel Laboratory of Neuroscience (FILN). Grants: ANR/CRCNS-BASCO, ANR-BALWM, ANR- BALAV1, France-Israel High Council for Science and Technology, LIA-FILN (CNRS), IRN-FICNC (CNRS).

## Appendix A: Correlations in binary networks

Here we calculate the equilibrium value and the stability of the quenched average correlations.

We define, for (*j, β*) ≠ (*i, α*), the out of equilibrium autoand crosscorrelations, 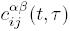 as

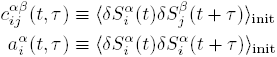

where ⟨ ⟩_init_ denotes averaging over many initial conditions choosen with a probability measure that, for simplicity, we choose such that 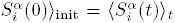. It is also convenient for the notation to define 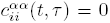. In this paper we focus on the equal time correlations, 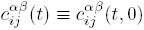.

For networks of binary neurons the dynamics of the equal-time crosscorrelations is given by [1, 28, 43]

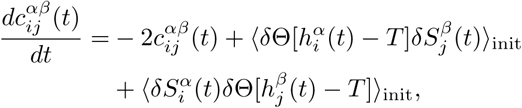

Where 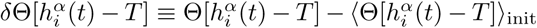.

If we make the *Ansatz* that correlations are weak, we can, to leading order, take 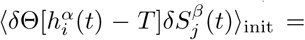 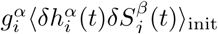, where 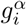 is the gain of neuron 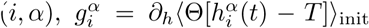, which is, to leading order, independent of the correlations [28, 43, 49].

Thus,

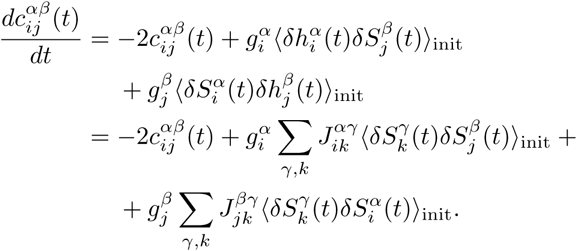

Using 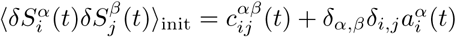 yields

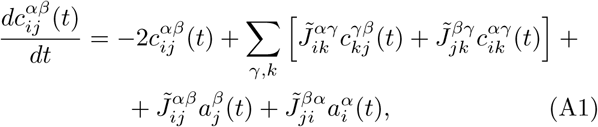

where 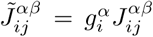. Note that, for weak correlations 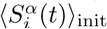 does not depend on the correlations so that, 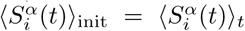 for our choice of initial conditions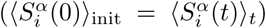. Hence, the equal time autocorrelation 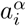 is independent of time and given by 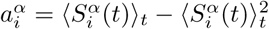.

We now average over the quenched disorder. Due to the rotational symmetry of the connection probabilities, 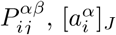 is a constant

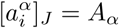

whereas 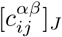 and 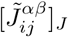 are functions of the difference in the location of neurons (*i, α*) and (*j, β*)

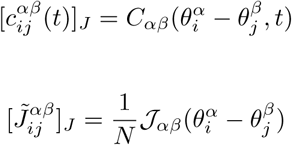

where 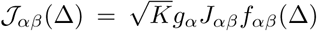. Here we have assumed that the correlations in the quenched disorder in the inputs to the neurons are small, such that, to leading order, the expected value of the gain does not depend on the neuronal position. We comment on this Ansatz in Appendix C.

Thus, for large *N*, the quenched average of Eq. (A1) yields

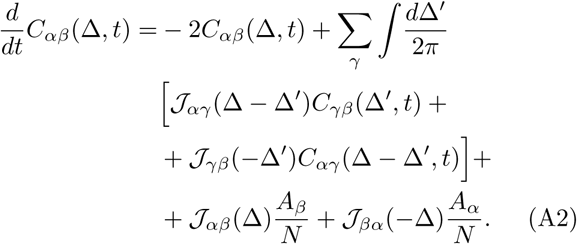

Here we have not taken into account that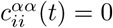. It is easy to see, however, that in the cross-correlations this neglects 𝒪 (1*/N* ^2^) corrections. We show below that these corrections are indeed negligeable in the large *N* limit.

The *n*^*th*^ Fourier mode of 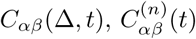 satisfies

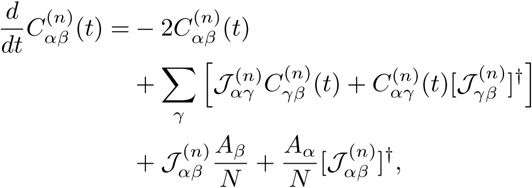

where 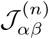 and 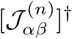 are the *n*^*th*^ Fourier mode of 𝒯 _*αβ*_(Δ) and 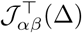, respectively.

This can be written more compactly as

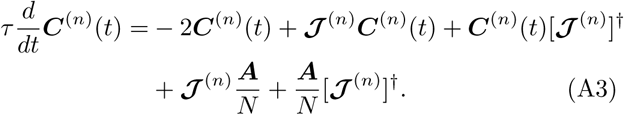

where ***X*** denotes the *D × D* matrix, *X*_*αβ*_, and ***A*** is the *D × D* matrix *A*_*αβ*_ *≡ δ*_*α,β*_*A*_*α*_. The equilibrium values of the Fourier components of equal-time correlation functions thus satisfy Eq. (26).

## Appendix B: Correlation theorems

In Appendix A we derived *N* ^2^*D*^2^ coupled linear differential equations that determine the evolution of equal-time cross-correlations for all neuronal pairs in the network. Averaging these equations over the quenched disorder and using the rotation invariance of the connection probabilities yields a set of *ND*^2^ coupled equations for the quenched averaged correlations. In Fourier space these equations lead to *N* independent sets of *D*^2^ coupled equations for the correlations. Here we prove Theorems 1 and 2 (see Section V) which state how these correlations scale with *N* and *K*.

To leading order, ***A*** is in<u>de</u>pendent of *K* and *N*, but 𝒯 ^(n)^ is proportional to 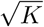. Accordingly, we define 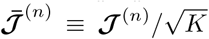 and 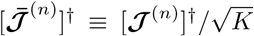. Thus, we can rewrite the evolution equation of the *n*^*th*^ mode of the correlations as

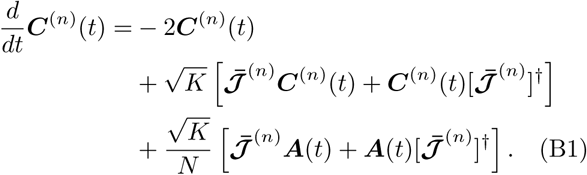

The *D × D* matrix, 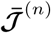can be written as

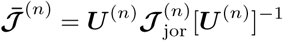

where 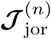 is the Jordan normal form of 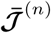 and [***U*** ^(n)^]^*-*^1^^ is the transformation matrix to the Jordan basis.

The matrix 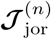 can be written as

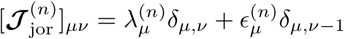

where 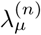 are the eigenvalues of 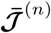 and 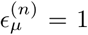 inside a Jordan block and is 0 otherwise (for clarity, in the Jordan basis we use the subscripts *μ* and *v*, rather than *α* and *β* that we use in the original basis).

Importantly, the Jordan form of a matrix and of its Hermitian conjugate are complex conjugate. We thus can write

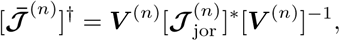

For notational convenience we will suppress the superscript (n)in the rest of this Appendix. Defining *Ĉ*as

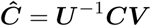

and inserting into Eq. (B1) yields

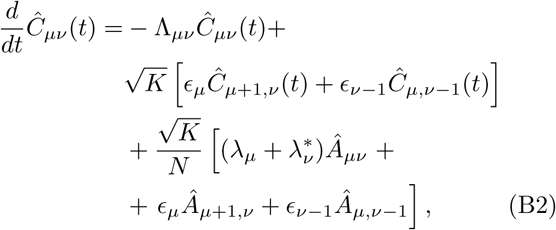

where we have defined 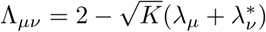 and

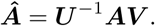

We now assume that the connectivity is such that the system is stable to perturbations of the locally averaged rates. This implies t<u>ha</u>t the real part of all eigenvalues of 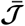 are less than 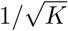. The real part of λ_*μv*_ is therefore positive and thus *Ĉ*^*(t)*^ converges to an equilibrium value, *Ĉ* ^∞^.

To see this, first consider *Ĉ*_*D,*1_*(t)* which satisfies

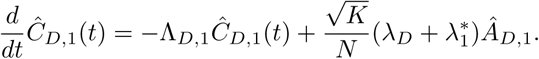

Therefore *Ĉ*_*D,*1_*(t)*, converges to its equilibrium value.

The evolution equations of *Ĉ*_*D,2*_can be written as

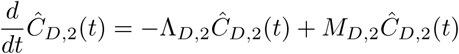

where *M*_*D,*2_ depends on *Ĉ*_*D,*1_*(t)* and *Ĉ*_*D,*1_*(t)*. Since *Â*_*D,*1_ and *Â*_*D,*2_ converges to its equilibrium value, *M*_*D,*2_ converges to a constant. Because Re(Λ_*D,2*_) >0, *Ĉ*_*D,*2_ also converges to its equilibrium value, 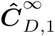. A similar argument shows that likewise *Ĉ*_*D-*1,1_ converges to 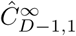. One then sees by recursion that the whole matrix *Ĉ* ∞ converges to its equilibrium value 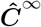. Hence, Eq. (26) determines the *stable* equilibrium values of the correlations.

From here we only consider the correlations at equilibrium and, for notational simplicity, we drop the superscript. *∞*

In the Jordan basis the equilibrium values of the correlations satisfy

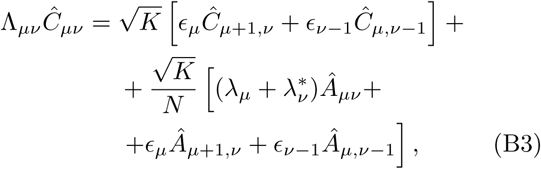

Let us now consider the case where 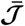 is diagonizable, so that *∈*_*μ*_ = 0 for all *μ*. In this case *Ĉ*_*μ v*_=0 for μ ≠v and

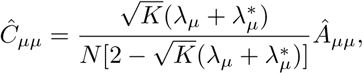

Thus

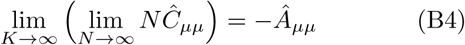

unless 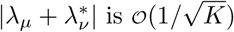, in which case

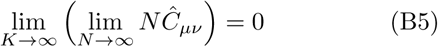

In the first situation *Ĉ* _*μ μ*_*= 𝒪*(1/*N)*. In the second situation *Ĉ* _*μ μ*_*= 𝒪*(1/*N)*.

Let us now assume that 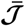 is not diagonizable. Then the *D × D* Jordan form of 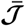 consists of *B* Jordan blocks (1*≤ B < D*) that we denote by 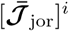*i* = 1;..., *B*. The size of the *i*th block will be denoted by *s*(*i*) *×s*(*i*). Without loss of generality, we can assume that the blocks are ordered in increasing size.

The indices *μ* and *v* of the elements of the Jordan block *i*, take values between *l*(*i*) and *h*(*i*), with 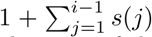 and 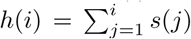. All the diagonal elements of this Jordan block are equal one of the eigen-values of 𝒯, which we denote by 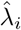. The off-diagonal elements, *∈*_*μ*_, are all equal to 1 except for *μ* = *h*(*i*) (*i* = 1,*.., B -* 1) for which *∈*_*μ*_ = 0.

The matrix *Ĉ* consists of *B*^2^ sectors that we denote by *S*_*ij*_. In the sector *S*_*ij*_, *μ ∈ {l*(*i*), *…, h*(*i*)*}* and *v∈ {l*(*j*), *…, h*(*j*) *}*.

Let us consider Eq. (B3) for *μ, v* in sector *S*_*ij*_. Since *∈*_*h*(*i*)_ and *∈*_*l*(*j*)*-*1_ are zero, the equation in this sector does not depend on elements of *Ĉ* outside of it. Thus, we can solve Eq. (B3) recursively to determine all the elements of 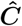 in this sector. The recursion goes as follows. First, one solves for 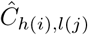

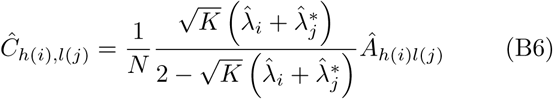

We can the solve Eq. (B3) to get *Ĉ* _*μv*_for μ, v = h(i) 1, l(j) and μ, v = h(i), l(j) + 1. This process can be repeated until all the elements of *Ĉ*in the sector S_*ij*_ are determined.

### 1. 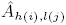are all non-zero

A similar recursion can be performed to estimate the order of magnitude of all the elements of *Ĉ*. First we obtain

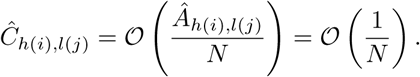

For 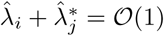 the recursion shows that we have

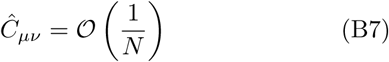

for all *μ, v* in sector *S*_*ij*_.

If 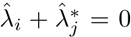, 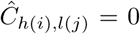 and by recursion all the other elements in sector *S*_*ij*_ are

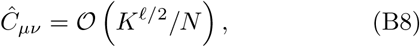

where *𝓁* = *v - μ* + *h*(*i*) *- l*(*j*). Thus, in this case *Ĉ* _*l*(*i*) *- h*(*j*)_ is the largest entry in the sector. It satisfies

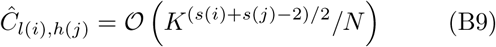

The scaling of the elements in sector *S*_*ij*_ thus depends 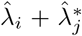. The stability of the dynamics imposes that 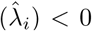 for all blocks (or positive but at most 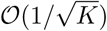, see next subsection) and thus the condition 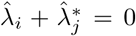 implies that, for large K, the real part of 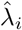 and 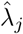 are zero. In other words, the real part of 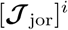 and 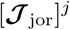 are shift matrices of size *s*(*i*)*×s*(*i*) and *s*(*j*) *× s*(*j*), respectively. For the sector *S*_*ii*_ it means that the real part of 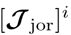 is a shift matrix of size *s*(*i*) *× s*(*i*).

Proving Correlation Theorem 1 is now straightforward. If the real part of all the *B* Jordan blocks of 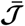 are different from a shift matrix, one finds that 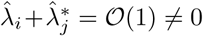 for all *i, j ∈{*1, *…, B}*. According to Eq. (B7), correlations are at most 𝒪 (1*/N*) in all of the *B*^2^ sectors of the matrix *Ĉ*. As a result, ***C***^(n)^= (1*/N*). On the other hand, if in each of the *B*^2^ sectors of the matrix ***C*** correlations are at most 𝒪 (1*/N*), this is also the case for the element of *Ĉ* in all the sectors. In this case, Eq. (B7) implies that there is no Jordan block in 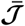 for which *Re*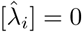.

Restoring the index *n* of the Fourier mode, one sees that if 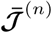 has at least one Jordan block whose real part is a shift matrix and denoting by *P* (n)the size of the largest shift, Eq. (B9) implies that 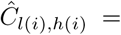 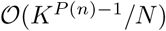, and thus also 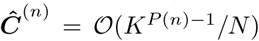. This proves Correlation Theorem 2.

According to Eq.(B9), the scaling of the correlation is 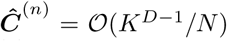 if and only if *P (n)*= *D*. When 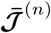 is a real matrix (when the probability of connections are symmetric in Δ), it means that the Jordan form of 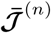 is a shift matrix of size *D*. This is equivalent for saying that 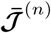 is nilpotent of degree *D*. On the other hand, if 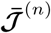 is a shift matrix of degree *D*, it has a Jordan form which is a shift matrix of degree *D*. According to Eq.(B9), the scaling of the correlation is then 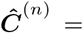 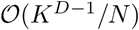. This proves the Corollary in section *V*.

When 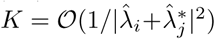 there is a crossover in sector *S*_*ij*_ of the matrix *Ĉ* between weak correlations (𝒪 (1*/N*)) and strong correlations (𝒪 (*K*^(*s*(*i*)+*s*(*j*)*-*^2^)*/*^2^^*/N*)). To see this, we note that if 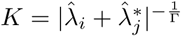, for Γ *≤* 1/2

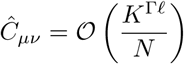

whereas for 1*/*2 *<* Γ *≤* 1

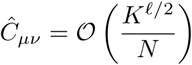

for *𝓁 ≠* 0 and

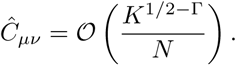

for *𝓁* = 0

### 2. Some elements of Â are zero

So far, we have assumed that all the elements in Â are non zero. The derivation can, however, be extended to include also situations where this is not the case as follows.

Elements Â _*μv*_=0 in a sector where 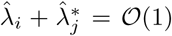 might result in some elements of 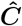in that sector to be equal to 0 rather than 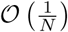. However, this will not affect the overall scaling of the correlations.

Elements Â _*μv*_=0in a sector Sij where 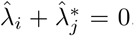, do not change the scaling in the matrix, provided that 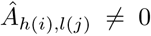. If 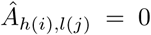 the effect depends on 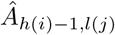 and 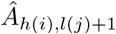 If at least one of them is nonzero, the order of the largest entry in the sector is decreased by 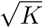. If both of them are zero, the order of this entry decreases at least by a factor of *K*. Reflecting on further element being zero, one reaches the following conclusion: Let *{μ*_*ij*_, *v*_*ij*_*}* be the indices *{μ, v}* in *S*_*ij*_ for which 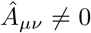, which maximize *μ - v*. Then, the maximal order of *Ĉ*_μv_ in the sector is 𝒪 (*K*^*P*^*i,j*^*-*^1^^*/N*), where

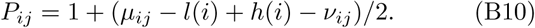

In conclusion: The highest order in *Ĉ* is *O*(*K*^*P-*^1^^*/N*), where *P* is the maximum value of the *P*_*ij*_ (*i, j ∈ {*1, *…, B}*) defined by: 1) *P*_*ij*_ = 1 for sector *S*_*ij*_ for which 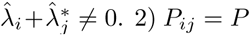 for sector *S*_*ij*_ in which 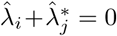 and all 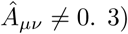 *P*_*ij*_ is given by Eq. (B10) if in sector 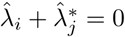 but some Â _μ_*v* are zero.

## Appendix C: Correlations of the quenched disorder

Let us define the correlation of the quenched disorder in the outputs of the neurons as

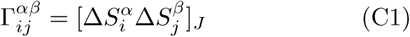

and the quenched disorder in the inputs,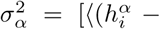 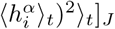, is

Finally, the gain of the neurons is

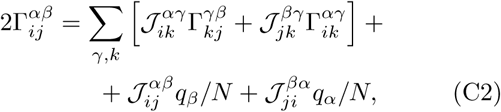

for (*i, α*) ≠ (*j, β*) and 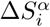 defined in Eq.(3). A derivation similar to that in Appendix A yields

Where 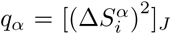. This equation can be solved using the same approach as in Appendices A-B. This analysis shows that correlations of the quenched disorder and correlations of the temporal fluctuations are of the same order. Thus, when the temporal fluctuations are small, the Ansatz in Appendix A, were we neglected the spatial fluctuations in the neuronal gain, is justified. If the correlations are too strong, the Ansatz may no longer be satisfied, but neither is the linearization assumed to derive equation (A2).

## Appendix D: Self-consistent equations for the autocorrelation and the gain

We follow the notations from [46]:

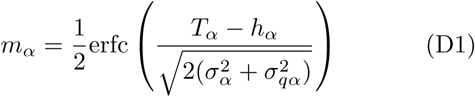

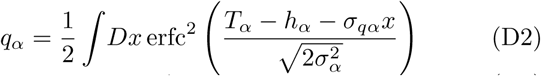

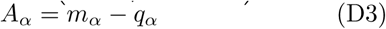

where the variance of the input noise, 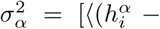 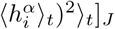,is

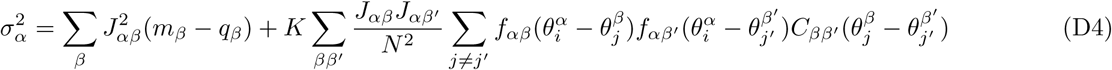

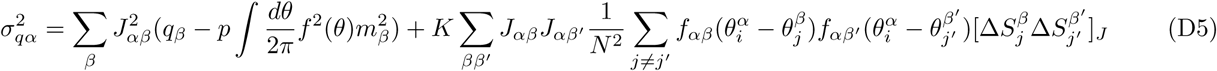

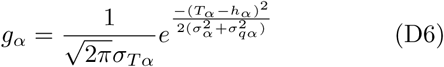

Equations (D1)-(D3) need to be solved self consistently. For simplicity, when we solved them we neglect 𝒪 (*p*) terms (but see [49]).

## Appendix E: Cross-correlations in two-population networks

For a two-population network Eq. (26) yields

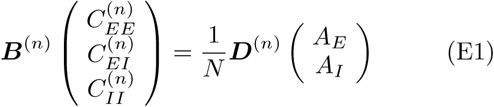

With 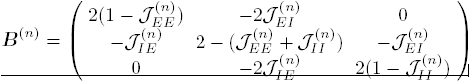 Solving this equation one gets

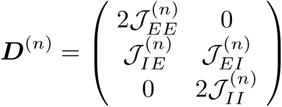

and

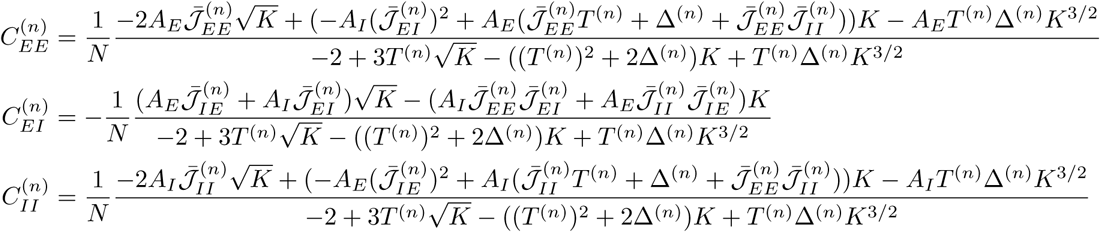

Where 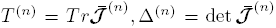. After some algebra, this equation can be rewritten as Eq. (27).

## References

[1] Roy J Glauber, “Time-dependent statistics of the ising model,” Journal of mathematical physics 4, 294–307 (1963).

[2] Mark C Cross and Pierre C Hohenberg, ““Pattern forma-tion outside of equilibrium,” Reviews of modern physics 65, 851 (1993).

[3] David Hansel and Haim Sompolinsky, “Chaos and syn-chrony in a model of a hypercolumn in visual cortex,” Journal of computational neuroscience 3, 7–34 (1996).

[4] Udo Seifert, “Stochastic thermodynamics, fluctuation theorems and molecular machines,” Reports on Progress in Physics 75, 126001 (2012).

[5] Fotis K Diakonos, AK Karlis, and P Schmelcher, “A universal mechanism for long-range cross-correlations,” EPL (Europhysics Letters) 105, 26004 (2014).

[6] Brent Doiron, Ashok Litwin-Kumar, Robert Rosenbaum, Gabriel K Ocker, and Krešimir Josić, “The mechanics of state-dependent neural correlations,” Nature neuro-science 19, 383–393 (2016).

[7] Tatjana Tchumatchenko, Aleksey Malyshev, Theo Geisel, Maxim Volgushev, and Fred Wolf, “Correlations and synchrony in threshold neuron models,” Physical Re-view Letters 104, 058102 (2010).

[8] Edwin M Maynard, Craig T Nordhausen, and Richard A Normann, “The utah intracortical electrode array: a recording structure for potential brain-computer inter-faces,” Electroencephalography and clinical neurophysi-ology 102, 228–239 (1997).

[9] Simon Peron, Tsai-Wen Chen, and Karel Svoboda, “Comprehensive imaging of cortical networks,” Current opinion in neurobiology 32, 115–123 (2015).

[10] Cyrille Rossant, Shabnam N Kadir, Dan FM Goodman, nJohn Schulman, Maximilian LD Hunter, Aman B Saleem, Andres Grosmark, Mariano Belluscio, George H Denfield, Alexander S Ecker, et al., Spike sorting for large, dense electrode arrays, Tech. Rep. (Nature Publishing Group, 2016).

[11] Ehud Zohary, Michael N Shadlen, and William T New-some, “Correlated neuronal discharge rate and its impli-cations for psychophysical performance,” (1994).

[12] LF Abbott and Peter Dayan, “The effect of correlated variability on the accuracy of a population code,” Neural computation 11, 91–101 (1999).

[13] Haim Sompolinsky, Hyoungsoo Yoon, Kukjin Kang, and Maoz Shamir, “Population coding in neuronal systems with correlated noise,” Physical Review E 64, 051904 (2001).

[14] Bruno B Averbeck, Peter E Latham, and Alexan-dre Pouget, “Neural correlations, population coding and computation,” Nature Reviews Neuroscience 7, 358–366 (2006).

[15] Gyorgy Buzsaki, Rhythms of the Brain (Oxford University Press, 2006).

[16] Charles M Gray, Peter K¨onig, Andreas K Engel, and Wolf Singer, “Oscillatory responses in cat visual cor-tex exhibit inter-columnar synchronization which reflects global stimulus properties,” Nature 338, 334–337 (1989).

[17] R Darshan, WE Wood, S Peters, A Leblois, and D Hansel, “A canonical neural mechanism for behavioral variability.” Nature communications 8, 15415 (2017).

[18] Neta Ravid Tannenbaum and Yoram Burak, “Shaping neural circuits by high order synaptic interactions,” PLoScomputational biology 12, e1005056 (2016).

[19] Donald Olding Hebb, The organization of behavior: A neuropsychological theory (Psychology Press, 2005).

[20] Guo-qiang Bi and Mu-ming Poo, “Synaptic modification by correlated activity: Hebb’s postulate revisited,” An-nual review of neuroscience 24, 139–166 (2001).

[21] Alexander S Ecker, Philipp Berens, Georgios A Keliris, Matthias Bethge, Nikos K Logothetis, and Andreas S Tolias, “Decorrelated neuronal firing in cortical micro-circuits,” science 327, 584–587 (2010).

[22] Marlene R Cohen and Adam Kohn, “Measuring and in-terpreting neuronal correlations,” Nature neuroscience 14, 811–819 (2011).

[23] Bryan J Hansen, Mircea I Chelaru, and Valentin Dragoi, “Correlated variability in laminar cortical circuits,” Neu-ron 76, 590–602 (2012).

[24] Maria C Dadarlat and Michael P Stryker, “Locomotion enhances neural encoding of visual stimuli in mouse v1,” Journal of Neuroscience 37, 3764–3775 (2017).

[25] Gideon Rothschild, Lior Cohen, Adi Mizrahi, and Israel Nelken, “Elevated correlations in neuronal ensembles of mouse auditory cortex following parturition,” Journal of Neuroscience 33, 12851–12861 (2013).

[26] Takaki Komiyama, Takashi R Sato, Daniel H OConnor, Ying-Xin Zhang, Daniel Huber, Bryan M Hooks, Mari-ano Gabitto, and Karel Svoboda, “Learning-related fine-scale specificity imaged in motor cortex circuits of behav-ing mice,” Nature 464, 1182–1186 (2010).

[27] James M Jeanne, Tatyana O Sharpee, and Timothy Q Gentner, “Associative learning enhances population cod-ing by inverting interneuronal correlation patterns,” Neu-ron 78, 352–363 (2013).

[28] Alfonso Renart, Jaime de la Rocha, Peter Bartho, Liad Hollender, Néstor Parga, Alex Reyes, and Kenneth D Harris, “The asynchronous state in cortical circuits,” science 327, 587–590 (2010).

[29] Timothy J Gawne and Barry J Richmond, “How inde-pendent are the messages carried by adjacent inferior temporal cortical neurons?” Journal of Neuroscience 13, 2758–2771 (1993).

[30] Timothy J Gawne, Troels W Kjaer, John A Hertz, and Barry J Richmond, “Adjacent visual cortical complex cells share about 20% of their stimulus-related informa-tion,” Cerebral Cortex 6, 482–489 (1996).

[31] Matthew A Smith and Adam Kohn, “Spatial and tem-poral scales of neuronal correlation in primary visual cortex,” The Journal of Neuroscience 28, 12591–12603 (2008).

[32] Diego A Gutnisky and Valentin Dragoi, “Adaptive coding of visual information in neural populations,” Nature 452, 220–224 (2008).

[33] Wyeth Bair, Ehud Zohary, and William T Newsome, “Correlated firing in macaque visual area mt: time scales and relationship to behavior,” Journal of Neuroscience 21, 1676–1697 (2001).

[34] Daeyeol Lee, Nicholas L Port, Wolfgang Kruse, and Apostolos P Georgopoulos, “Variability and correlated noise in the discharge of neurons in motor and parietal areas of the primate cortex,” The Journal of neuroscience 18, 1161–1170 (1998).

[35] Robert B Levy and Alex D Reyes, “Spatial profile of excitatory and inhibitory synaptic connectivity in mouseprimary auditory cortex,” The Journal of Neuroscience 32, 5609–5619 (2012).

[36] Elodie Fino and Rafael Yuste, “Dense inhibitory connec-tivity in neocortex,” Neuron 69, 1188–1203 (2011).

[37] Hiroki Tanaka, Hiroshi Tamura, and Izumi Ohzawa, “Spatial range and laminar structures of neuronal cor-relations in the cat primary visual cortex,” Journal of neurophysiology 112, 705–718 (2014).

[38] Shervin Safavi, Abhilash Dwarakanath, Vishal Kapoor, Joachim Werner, Nicholas G Hatsopoulos, Nikos K Lo-gothetis, and Theofanis I Panagiotaropoulos, “Non-monotonic spatial structure of interneuronal correlations in prefrontal microcircuits,” bioRxiv, 128249 (2017).

[39] Robert Rosenbaum, Matthew A Smith, Adam Kohn, Jonathan E Rubin, and Brent Doiron, “The spatial structure of correlated neuronal variability,” Nature Neu-roscience 20, 107–114 (2017).

[40] Daniel J Denman and Diego Contreras, “The structure of pairwise correlation in mouse primary visual cortex reveals functional organization in the absence of an ori-entation map,” Cerebral Cortex 24, 2707–2720 (2014).

[41] Tom Binzegger, Rodney J Douglas, and Kevan AC Mar-tin, “A quantitative map of the circuit of cat primary visual cortex,” Journal of Neuroscience 24, 8441–8453 (2004).

[42] Moshe Abeles, Corticonics: Neural circuits of the cerebral cortex (Cambridge University Press, 1991).

[43] Iris Ginzburg and Haim Sompolinsky, “Theory of corre-lations in stochastic neural networks,” Physical review E 50, 3171 (1994).

[44] Jérémie Barral and Alex D Reyes, “Synaptic scaling rule preserves excitatory-inhibitory balance and salient neu-ronal network dynamics,” Nature neuroscience 19, 1690–1696 (2016).

[45] Carl van Vreeswijk and Haim Sompolinsky, “Chaos in neuronal networks with balanced excitatory and in-hibitory activity,” Science 274, 1724–1726 (1996).

[46] Carl van Vreeswijk and Haim Sompolinsky, “Chaotic bal-anced state in a model of cortical circuits,” Neural com-putation 10, 1321–1371 (1998).

[47] Michael Monteforte and Fred Wolf, “Dynamic flux tubes form reservoirs of stability in neuronal circuits,” Physical Review X 2, 041007 (2012).

[48] Alex Roxin, Nicolas Brunel, David Hansel, Gianluigi Mongillo, and Carl van Vreeswijk, “On the distribution of firing rates in networks of cortical neurons,” The Journal of neuroscience 31, 16217–16226 (2011).

[49] Moritz Helias, Tom Tetzlaff, and Markus Diesmann, “The correlation structure of local neuronal networks in-trinsically results from recurrent dynamics,” PLoS Com-put Biol 10, e1003428 (2014).

[50] David Hansel and Haim Sompolinsky, “Synchronization and computation in a chaotic neural network,” Physical Review Letters 68, 718 (1992).

[51] Zoltán F Kisvárday, E Toth, Martin Rausch, and Ulf T Eysel, “Orientation-specific relationship between popula-tions of excitatory and inhibitory lateral connections in the visual cortex of the cat.” Cerebral cortex (New York, NY: 1991) 7, 605–618 (1997).

[52] Armen Stepanyants, Luis M Martinez, Alex S Ferecsk´o, and Zoltán F Kisvárday, “The fractions of short-and long-range connections in the visual cortex,” Proceedings of the National Academy of Sciences 106, 3555–3560 (2009).

[53] Rani Ben-Yishai, David Hansel, and Haim Sompolinsky, “Traveling waves and the processing of weakly tuned in-puts in a cortical network module,” Journal of computa-tional neuroscience 4, 57–77(1997).

[54] David Hansel and Haim Sompolinsky, “13 modeling fea-ture selectivity in local cortical circuits,” (1998).

[55] C Van Vreeswijk and H Sompolinsky, “Irregular activity in large networks of neurons,” Methods and models in neurophysics. Amsterdam: Elsevier (2005).

[56] Robert Rosenbaum and Brent Doiron, “Balanced net-works of spiking neurons with spatially dependent recur-rent connections,” Physical Review X 4, 021039 (2014).

[57] Eric Shea-Brown, Krešimir Josić, Jaime de La Rocha, and Brent Doiron, “Correlation and synchrony transfer in integrate-and-fire neurons: basic properties and conse-quences for coding,” Physical review letters 100, 108102 (2008).

[58] Jaime De La Rocha, Brent Doiron, Eric Shea-Brown, Krešimir Josić, and Alex Reyes, “Correlation between neural spike trains increases with firing rate,” Nature 448, 802–806 (2007).

[59] Vladimir Litvak, Haim Sompolinsky, Idan Segev, and Moshe Abeles, “On the transmission of rate code in long feedforward networks with excitatory–inhibitory balance,” The Journal of neuroscience 23, 3006–3015 (2003).

[60] Nelson Cortes and Carl van Vreeswijk, “Pulvinar thala-mic nucleus allows for asynchronous spike propagation through the cortex,” Frontiers in computational neuro-science 9 (2015).

[61] Matthew A Smith, Xiaoxuan Jia, Amin Zandvakili, and Adam Kohn, “Laminar dependence of neuronal correla-tions in visual cortex,” Journal of neurophysiology 109, 940–947 (2013).

[62] Nicolas Brunel, “Dynamics of sparsely connected net-works of excitatory and inhibitory spiking neurons,” Journal of computational neuroscience 8, 183–208 (2000).

[63] Kanaka Rajan and LF Abbott, “Eigenvalue spectra of random matrices for neural networks,” Physical review letters 97, 188104 (2006).

